# An Abundance of Free Proteasomal Regulatory (19S) Particles Regulate Neuronal Synapses Independent of the Proteasome

**DOI:** 10.1101/2022.10.17.512557

**Authors:** Chao Sun, Kristina Desch, Belquis Nassim-Assir, Stefano L. Giandomenico, Paulina Nemcova, Julian D. Langer, Erin M. Schuman

## Abstract

The major protein-degradation machine, the proteasome, functions at brain synapses and regulates long-term information storage. Here we found that the two essential subcomplexes of the proteasome, the regulatory (19S) and catalytic (20S) particles that recognize and degrade substrates, are differentially distributed within individual rat cortical neurons. Using superresolution microscopy, we discovered a surprising abundance of free particles (19S) near synapses. The free neuronal 19S particles bind and deubiquitylate Lys63-ubiquitin, a non-proteasome targeting ub-linkage. Pull-down assays revealed a significant over-representation of synaptic molecules as Lys63 interactors. Inhibition of 19S deubiquitylase activity significantly altered excitatory synaptic transmission and reduced the synaptic availability of AMPA receptors at multiple trafficking points in a proteasome-independent manner. Together, these results reveal a moonlighting function of the regulatory proteasomal subcomplex near synapses.

**One-Sentence Summary:** The 19S subcomplex of the major protein-degradation machine, the proteasome, functions without its 20S partner to populate and regulate synapses.

## Introduction

An individual neuron hosts about 10^4^ synapses in its expansive dendritic and axonal arbor (*1*). To support parallel synaptic computation and plasticity, both protein synthesis and degradation machines are localized to remodel individual synaptic proteomes on site (*2–5*). Ribosomes and proteasomes are assembled from multiple functional subunits, giving rise to different assembly states near synapses (*6–9*). The different assembly states can exhibit distinct, novel functions within neuronal subcellular compartments. For example, while polyribosomes (multiple ribosomes engaged with an mRNA) have been regarded as the main form of protein-synthesis machine (*10*), single ribosomes (monosomes) have recently been described as an important source of translation near synapses (*6*). Data like these raise the possibility that synapses can use alternative machine formats to carry out different functions, akin to protein ‘moonlighting’. However, so far little is known about the potential synaptic adaptation of multiprotein complexes such as their local assembly states and potentially unique functions.

The proteasome is a multiprotein complex responsible for most cytosolic protein degradation in cells (*11*); it operates at synapses (*5*, *12*) and its function is necessary for long-term memory (*13, 14*). Proteasomes consist of regulatory (19S) and catalytic particles (20S) (*15*). Proteins are targeted for proteasomal degradation by the attachment of ubiquitin chains that are primarily linked at Lysine-48 (Lys48) (*16*). There exists, however, another major type of ubiquitylation in mammalian cells, ubiquitin chains linked via Lysine-63 (Lys63-ubiquitylation) (*17*); Lys63-ubiquitylated proteins are not targeted for proteasomal degradation (*18*). The process of proteasomal degradation includes the Lys48-deubiquitylation of protein targets by a deubiquitylase in the 19S particle followed by immediate protease-mediated degradation within the barrel-shaped 20S particle (*15*). Thus, canonical proteasome function requires the stable assembly of the regulatory 19S and the catalytic 20S particles (*19*, *20*). As might be expected, the average expression levels of 19S and 20S protein subunits are also stoichiometric for singly-capped proteasomes (26S), consisting of one 19S particle and one 20S particle (*6*, *21*). Consistent with this subunit expression, an electron cryo-tomography study detected assembled neuronal proteasomes primarily as singly-capped 26S proteasomes with a minor fraction of 30S doubly-capped proteasomes (*7*). In addition, neuronal synapses sequester both types of proteasomal particles (19S and 20S) in an activity-dependent manner (*5*).

The average half-life of the proteins that make up the neuronal 19S particle, however, is significantly different from those that comprise the 20S particle, suggesting that the two subcomplexes do not turn over in concert (*22*). In addition, emerging evidence also points to distinct functions of free 20S particles (*23*, *24*), whose detection remains elusive in electron cryo-tomography. These data suggest that the 26S proteasome is not the only assembly or functional state of synaptic proteasome particles. Considering the huge protein turnover demand in neuronal processes (*1*, *25*), the nature of synaptic proteasomal assembly states represents a major knowledge gap. Here we show that neuronal proteasomal regulatory and catalytic particles are unevenly distributed within neurons both *in vivo* and *in vitro*. An ‘excess’ of proteasomal regulatory particles 19S (relative to 20S) was detected near synapses. Combining single-molecule localization microscopy and native protein gel electrophoresis, we profiled the dendritic proteasome assembly states and reveal a moonlighting function for free 19S proteasomal regulatory particles as deubiquitylases (DUBs) against Lys63 ubiquitin chains which target synaptic proteins. Excess proteasomal regulatory particles thus modify an unconventional form of ubiquitylation and regulate crucial synaptic proteins in parallel to their more conventional role in proteasome degradation.

## Results

To investigate the subcellular distribution of proteasomal regulatory 19S particles and catalytic 20S particles, we conducted immunolabelling for 19S and 20S protein subunits in mouse brain slices and in cultured rat cortical neurons using confocal microscopy. Because the 20S particle contains two copies of each constituent protein subunit while the 19S particle contains only one copy of each protein subunit (Fig. 1A) (*15*), we targeted the two copies of PSMA7 subunit in each 20S particle, and two subunits of each 19S particle (PSMC1 and PSMD6) using validated antibodies (Fig. S1). If the 26S proteasome is the main assembly state of both neuronal proteasomal particles (Fig. 1A), then the 19S and 20S immunofluorescence should distribute in a consistent ratio in neuronal cell bodies and their processes. However, this was not what we observed. In coronal mouse brain slices, the relative fluorescence of the 19S and 20S subunits varied greatly across brain regions (Fig. 1B). In hippocampal area CA1 where neuronal cell bodies and processes exhibit a clear laminar organization (Fig. 1B right inset), the 20S fluorescence was relatively enriched in the somata layer (stratum pyramidale) while the 19S fluorescence was enriched in the neuropil layer (stratum radiatum). This subcellular heterogeneity of 19S/20S distribution was detected with greater resolution in cultured rat cortical neurons. As observed *in vivo*, the 19S subunit was significantly more abundant in dendrites and axons whereas the 20S subunit was more abundant in the cell body (Fig. 1C&D). Antibodies targeting different proteasome protein constituents of the 19 and 20S subcomplexes generated similar distribution patterns in cultured rat hippocampal neurons (Fig. S1C). These observations indicate that the 19S and 20S proteasome subcomplexes are differentially distributed in neuronal subcellular compartments, raising the possibility of different proteasome formats at synapses.

**Fig. 1.**
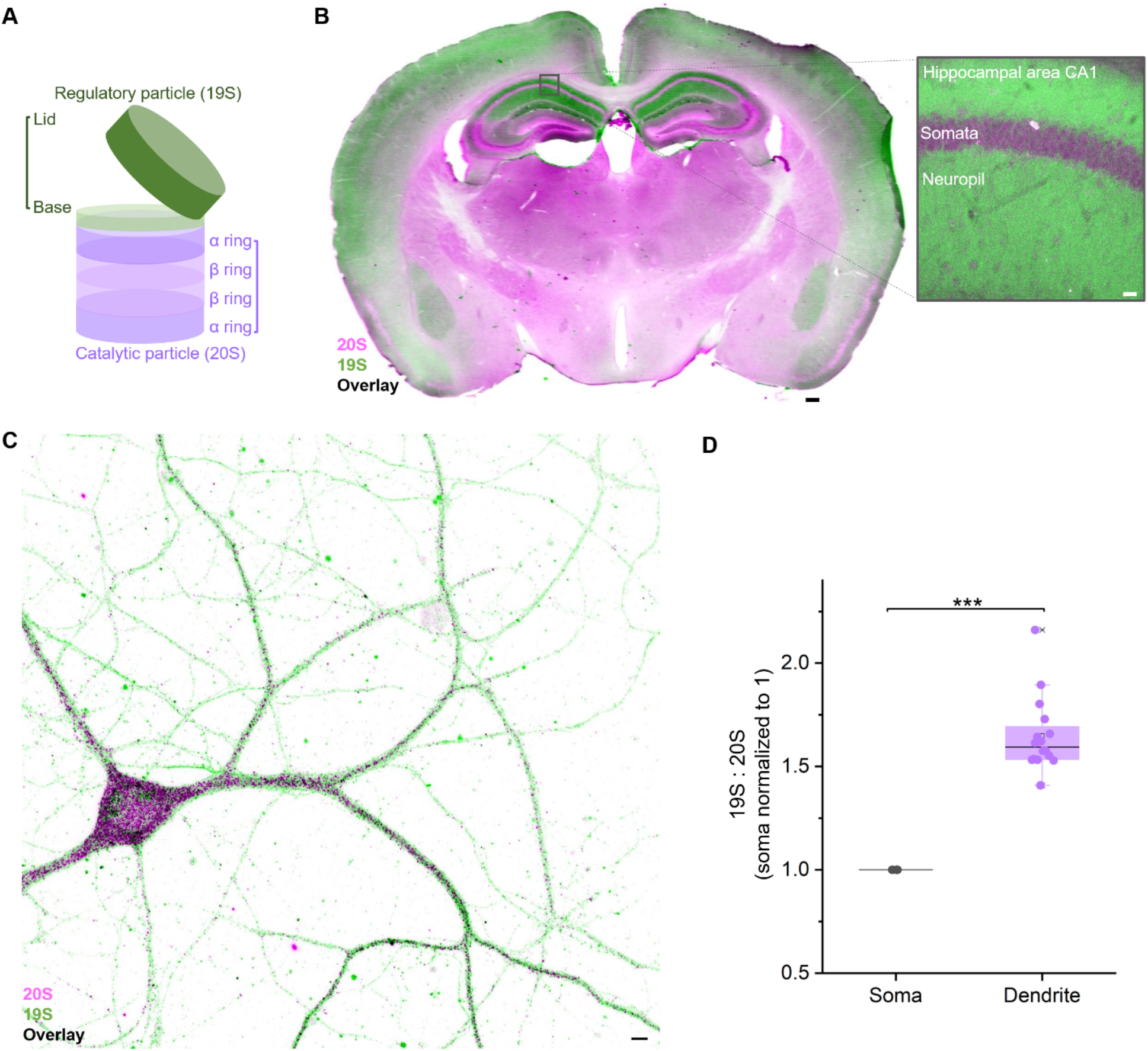
Proteasomal regulatory (19S) and catalytic (20S) particles are differentially distributed in neuronal subcellular compartments. **(A)** Illustration of a singly-capped proteasome (26S), where each 26S contains one copy of each 19S subunit (green) and two copies of each 20S subunit (magenta). **(B)** Immunofluorescence staining of a coronal section of the mouse brain, with 19S in green and 20S in magenta. Scale bar: 250 μm. Right inset is a confocal micrograph of the hippocampal area CA1 showing the enriched 19S staining (green) in the stratum radiatum (neuropil) and enriched 20S staining (magenta) in the stratum pyramidale (somata). Scale bar: 20 μm. **(C)** Confocal micrograph of a mature rat primary cortical neuron immunolabelled for the 19S (green) and 20S (magenta) particles. Scale bar: 4 μm. **(D)** Scatter plot showing a significant higher ratio of 19S: 20S particle fluorescence intensity in dendrites (17 dendritic branches) compared to in soma (normalized to 1.0; 6 cell bodies). *** = p <0.001 via a two-sample T test.

To elucidate further the abundance and assembly states of 19S and 20S near synapses, we visualized dendritic 19S and 20S particles using quantitative, multiplexed, single-molecule localization microscopy (DNA-PAINT; see methods) (*4, 26*). The 19S and 20S particles were detected as individual clusters of localizations at single-molecule resolution. We detected assembled proteasomes as the colocalization between 19S and 20S particles (Fig. 2A; also see method). On average, 2.5 copies of assembled (19S + 20S) proteasome particles were found per μm dendrite (Fig. 2B), with ~80% of the detected assembled proteasomes containing one 19S and one 20S (singly-capped proteasome 26S), and ~20% containing one 20S and two 19S copies (doubly-capped proteasome 30S). The detected ratio between 26S and 30S proteasomes was similar to what was observed by a recent *in situ* electron cryo-tomography study in neurons (*7*).While the 26S was the main form of assembled proteasome detected, free 19S and 20S subcomplexes were also abundant in neuronal dendrites (Fig. 2C). The ratio of total (free + assembled) 19S to 20S particles in dendrites was ~2:1 (Fig. 2D), exceeding the expected 1:1 ratio needed to assemble the 26S proteasome. Despite the super-stoichiometric presence of the 19S in dendrites, nearly half of the 20S particles remained uncapped (Fig. 2E). Correspondingly, ~70% of the 19S particles were detected in isolation (“free”) (Fig. 2F). While the free 20S proteasome has been implicated as an independent degradation machine with distinct protein substrates (*23*),the abundance of free 19S particles near synapses has not been previously detected.

**Fig. 2.**
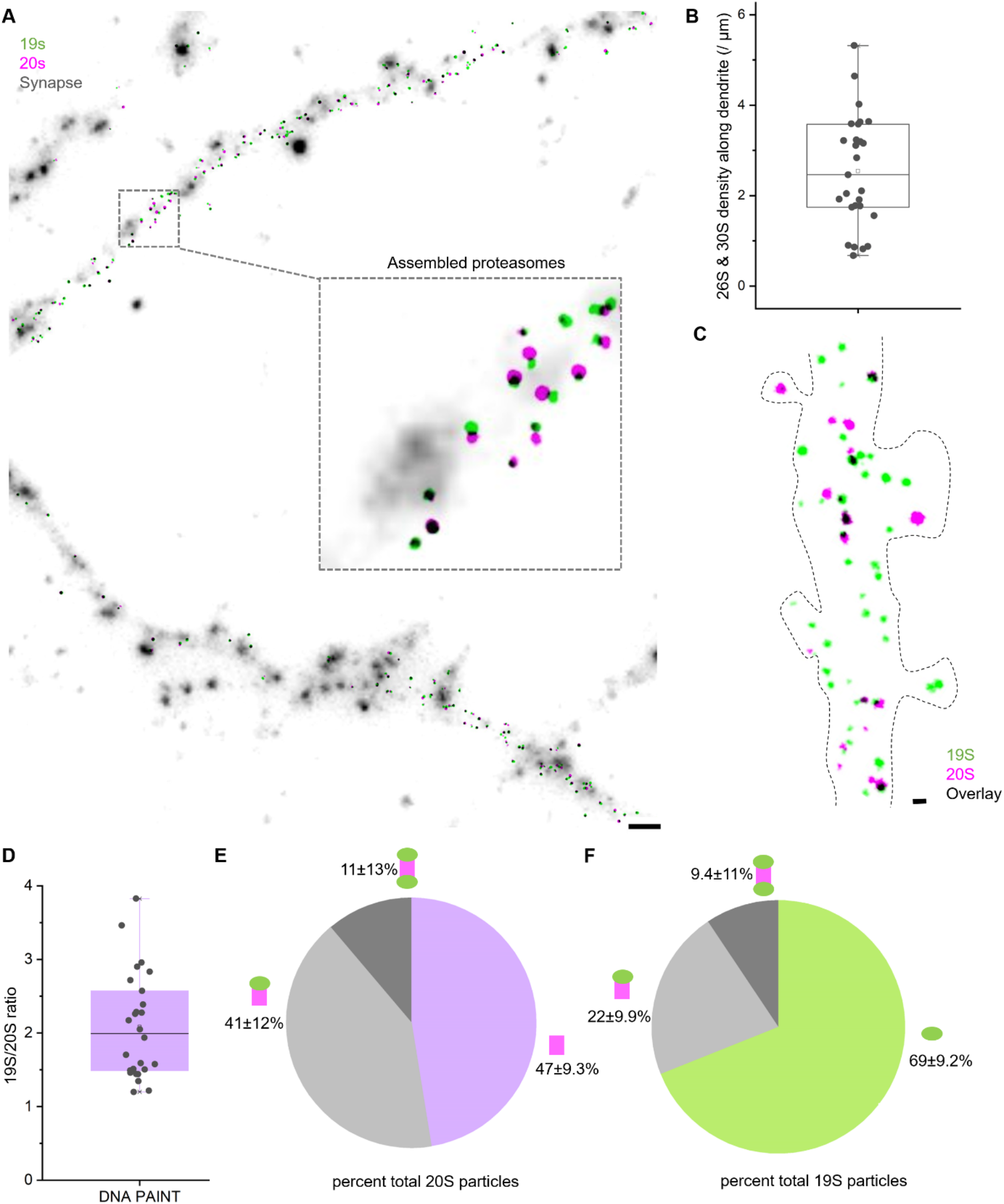
The assembly states of proteasome 19S and 20S proteasomes in neuronal dendrites. **(A)** Single-molecule localization micrograph of super-resolved 26S and 30S proteasomes in neuronal dendrites and synapses (grey; immunostaining against Bassoon) via DNA PAINT, with 19S in green, 20S in magenta, and overlay in black. Scale bar: 2 μm. Inset shows an enlarged region containing multiple proteasomes as paired magenta and green puncta. **(B)** A scatter plot quantifying the abundance of 26S and 30S proteasomes along 27 dendrites from 6 neurons, with an average density of 2.5 ± 1.2 copies per μm dendrite. **(C)** shows a dendritic segment with superresolved 19S particle (green), 20S particle (magenta), as well as their colocalization in 26S and/or 30S proteasomes (in black). **(D)** A scatter plot showing the ratio of 19S particles vs. 20S particles in 27 dendrites from 6 neurons obtained by DNA PAINT, with an average value (1.9 ± 0.5) greater than 1. **(E)** Pie chart showing the percentage of dendritic 20S particles in solo, singly-capped (26S), and doubly-capped (30S) states. **(F)** Pie chart showing the percentage of dendritic 19S particles in solo, singly-capped (26S), and doubly-capped (30S) states. The average values were obtained from 27 dendrites, 6 neurons.

We considered whether neurons possess elevated 19S abundance near synapses to increase the likelihood of capping 20S particles (thus generating 26S and 30S proteasomes). To test this, we harvested somata- and neurite-enriched neuronal lysates from cortical neurons cultured in compartmentalized chambers (*8*) (Fig. 3A; see also Fig. S1D). Using native protein gel electrophoresis, neuronal proteasomes comprising different assembly states (20S, 26S, and 30S) were separated as three distinct bands that were visualized by immunoblotting against the 20S subunit PSMA7 (Fig. 3B) (*27*). In the neurite fraction, we found that ~50% of the total 20S immunofluorescence intensity was present in the free 20S band of neurites (Fig. 3C), consistent with our single-molecule localization data (Fig. 2E). Approximately the same fraction of total 20S was present in the free 20S band from the somata-enriched fraction (Fig. 3C). As such, the super-stoichiometric presence of 19S in dendrites and synapses does not elevate the percentage of capped 20S proteasomal particles (26S or 30S) in this same compartment. These data suggest that the free 19S particles may carry out an, as yet, unknown, proteasome-independent function at synapses.

**Fig. 3.**
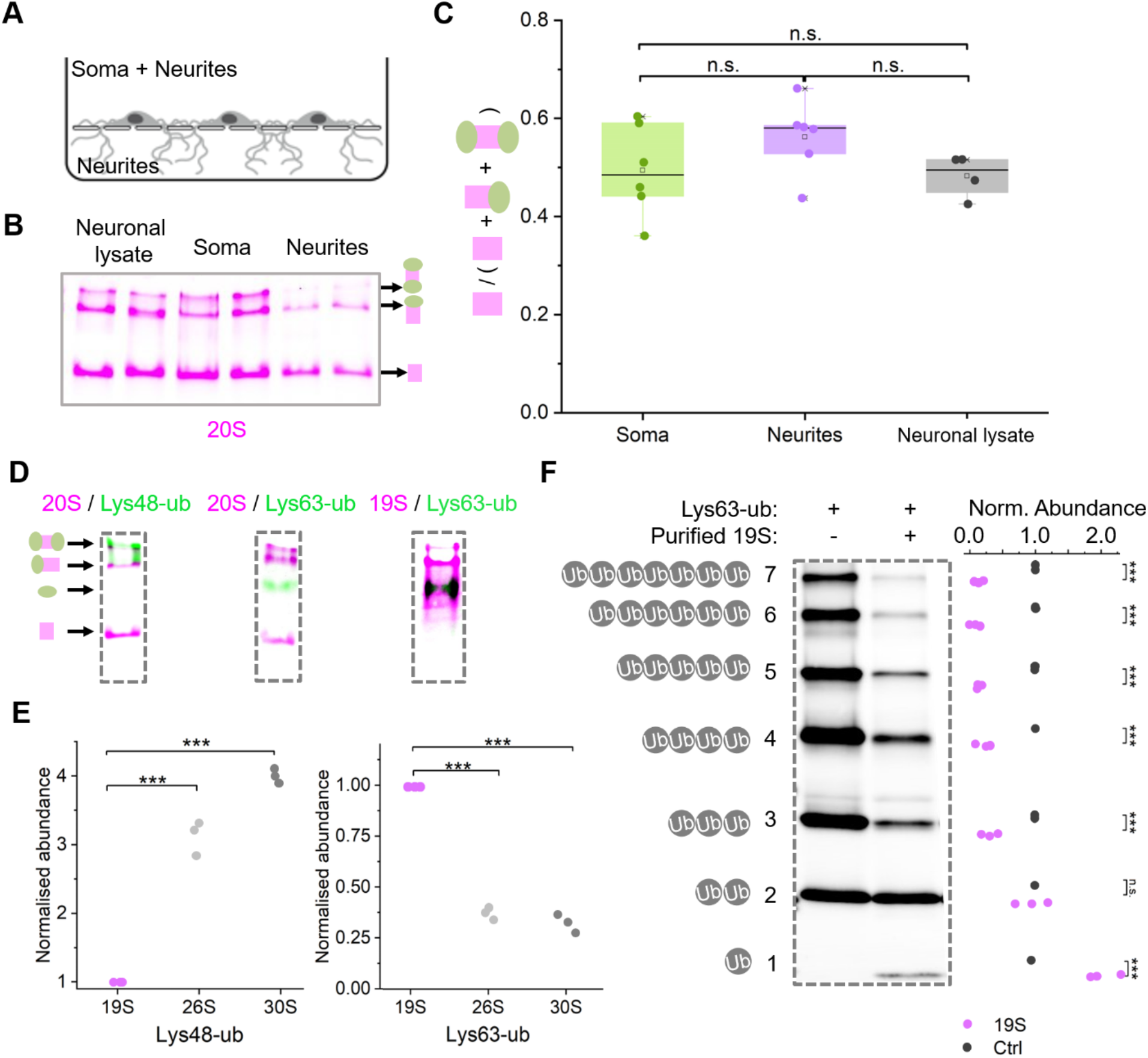
Free 19S proteasome regulatory particles function as proteasome-independent deubiquitylases acting on Lys63-ubiquitin. **(A)** Scheme of a compartmentalized chamber where cultured neurons can extend their neurites (dendrites and axons) through the porous membranes into the bottom compartment while their cell bodies (and some processes) remain in the upper compartment. **(B)** Native protein gel electrophoresis of neuronal lysate, soma-enriched lysate, and neurites-enriched lysate immunolabelled against the 20S particle (magenta; anti-PSMA7). The three bands, starting from the top, are the 30S, 26S, and free 20S proteasomes, with the respective cartoons indicating the capped (green + magenta) and uncapped states (magenta) on the right. **(C)** Box plots showing the fractions of free 20S obtained from native protein gel electrophoresis of the soma-enriched, neurites-enriched, and neuronal lysates. An analysis of variance (ANOVA) with post hoc Bonferroni test detected no significant differences in the free 20S between the compartments. **(D)** Native protein gel electrophoresis showing the differential association of the 19S to linkage-specific ubiquitylation. Left shows the predominant association of lys48- ubiquitylation with 19S bound to 20S (26S and 30S; anti-PSMA7 and anti-Lys48-ub). Middle shows the predominant presence of lys63-ubiquitylation at solo 19S band (anti-PSMA7 and anti-Lys63-ub). Right shows the predominant association of lys63-ubiquitylation with free 19S band (anti-PSMC1/PSMD6/Uchl5 followed by re-probing with anti-Lys63-ub). Note that free 19S bands appears above solo 20S band. Cartoons on the left shows 30S, 26S, 19S, and 20S from top to bottom. **(E)** scatter plots showing the normalized ubiquitin intensities associated with free 19S, 26S, and 30S particles in native protein gel electrophoresis. ANOVA with post-hoc Bonferroni test detected significantly less Lys48-ubiquitylation and significantly more Lys63-ubiquitylation in free 19S particles when comparted to 26S and 30S particles. *** = p< 0.001. **(F)** SDS-PAGE showing deubiquitylation of lys63-polyubiquitin chains by the free 19S particle following 2 hr coincubation. Scatter plots show the normalized band intensities for different ubiquitin chain lengths in control (dark grey) and 19S (magenta) lanes. Two-sample T-tests detected a significant increase in mono ubiquitin and significant decreases in the higher-molecular weight ubiquitin chains (3-7 repeats) following co-incubation with purified 19S particles (*** = p <0.001).

While the 19S regulatory particle does not degrade protein substrates, it is known to cleave ubiquitin chains (deubiquitylate, DUB) from ubiquitylated proteins at an early step in proteasome processivity (*28*). This 19S DUB activity is essential for subsequent protein degradation by the 20S particle as well as ubiquitin recycling. We thus hypothesized that the free 19S particles may operate as independent DUBs to reverse protein ubiquitylation, reasoning that this activity could serve to suppress proteasome degradation near synapses. We first examined the type of ubiquitin chain that is associated with the free 19S subunit (and the other proteasome assembly states) using native protein gel electrophoresis and antibodies that recognize Lysine 48-linked ubiquitin chains (Lys48-ub) or Lysine 63-linked ubiquitin chains (Lys63-ub). As previously reported, we found that Lys48-ub was mostly bound to 26S and 30S proteasomes (Fig. 3D), but not to the free 19S particle (*27*). This indicates that the free 19S particle does not interact (to an appreciable extent) with Lys48-ubiquitylated proteins in cell lysates (Fig. 3E). In contrast, we found that the Lys63-ubiquitylated proteins were predominantly associated with the free 19S particle rather than with the 26S or 30S particles (Fig. 3D & 3E). To test directly whether Lys63-ubiquitin chains undergo deubiquitylation as a result of their association with the free 19S particle, we reconstituted an enzymatic reaction that included purified free 19S particles and Lys63-polyubiquitin chains *in vitro*. When the 19S particle was added to the purified Lys63-ubiquitin chain, we observed the emergence of mono-ubiquitin products due to 19S deubiquitylation, as well as the loss of higher molecular-weight polyubiquitin species (Fig. 3F). To test whether this DUB activity is specific for the free 19S particle, we conducted the same experiment with the purified 26S proteasome and found it did not deubiquitylate Lys63-ub chains to the same extent under the same conditions (Fig. S2). These data indicate that the free 19S particle can act as a proteasome-independent DUB against Lys63 ubiquitylation. Given the abundance of the free 19S near synapses, these data raise the possibility that Lys63-ub-modified synaptic proteins could be regulated by the free 19S activity.

To investigate the potential neuronal substrates of Lys63-ubiquitylation, we performed immunoprecipitation of Lys63-ubiquitylated proteins using anti-Lys63-ub tandem ubiquitin binding entities (TUBEs) in cortical neuron lysates (see methods) and subsequent mass spectrometry-based proteomic analysis (Fig. S3 & S4). We compared the Lys63-ubiquitylated substrates to those identified by immunoprecipitation (IP) of Lys48-ubiquitylation as well as pan-ubiquitylation (Ub; with unspecified linkages). Amongst the > 2000 proteins identified in the pull-down samples, 650 protein species in the Lys63-ub IP were unique or enriched (compared to a “beads only” pull-down background), while 430 or 1841 unique/enriched proteins were detected in the Lys48-ub or pan-ubiquitin IP, respectively (Fig. 4A). Consistent with previous studies, several ribosomal proteins were Lys63-ubiquitylated (*29*). However, in addition, the Lys63-ub IP contained many synaptic proteins (Fig. 4A), including essential presynaptic proteins (e.g. syntaxin, synaptogyrin, and synaptotagmin) and neurotransmitter receptors and ion channels (AMPA receptors, voltage-gated sodium channels, calcium channels, and potassium channels) (See Data S1). In contrast, immunoprecipitation of Lys48-ubiquitylated proteins mainly yielded hits that enriched for proteasomal components, consistent with its predominant association with the 26S & 30S proteasomes in native protein gel electrophoresis (Fig. 3D). As such, these results suggest that the free 19S may act as a DUB for Lys63-ub modified synaptic proteins.

**Fig. 4.**
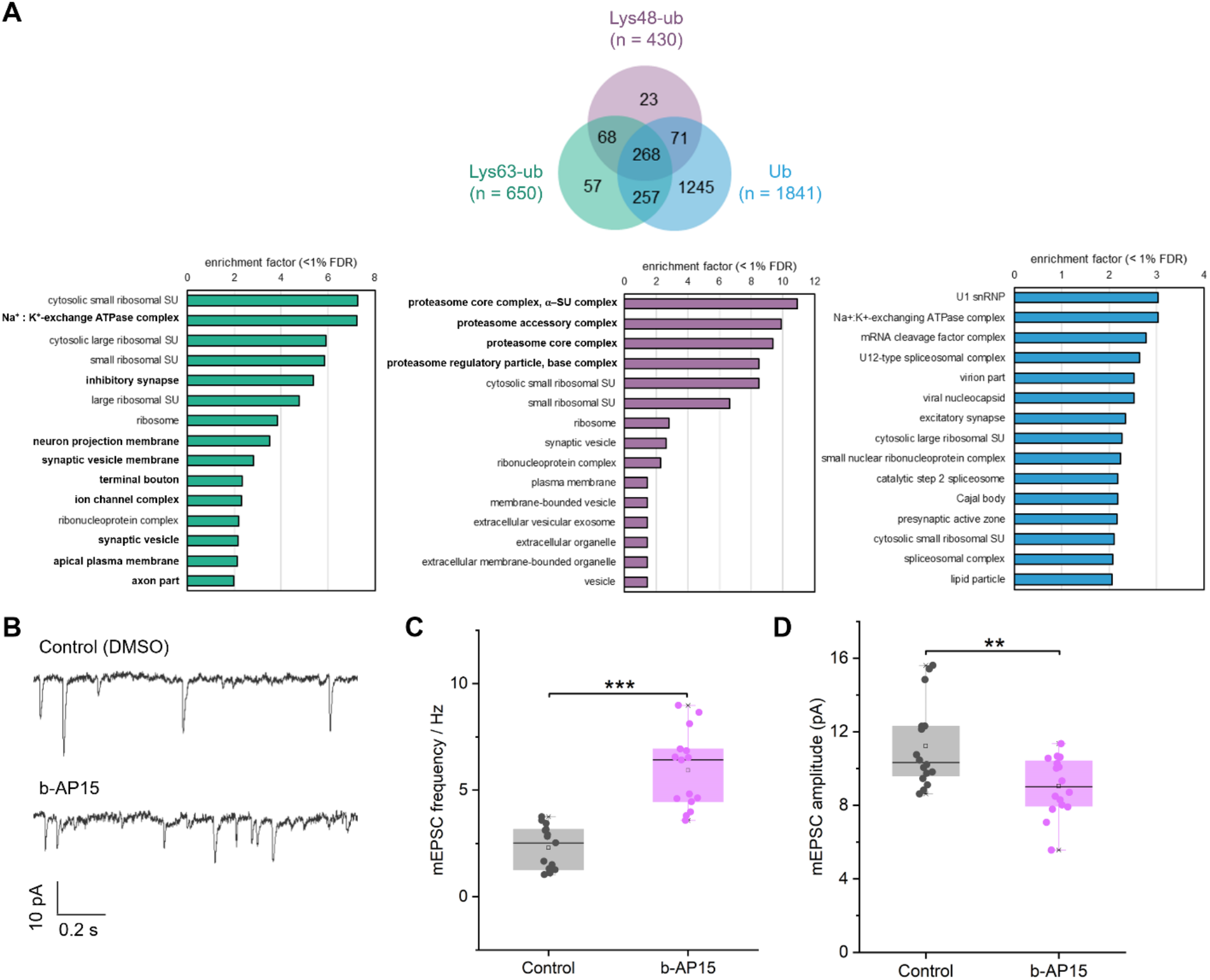
Synaptic proteins are major interactors of Lys63-ubiquitylation. **(A)** Top 15 gene-ontology cellular component terms enriched in Lys63-ub (left; based on 650 out of 2229 proteins), Lys48-ub (middle; based on 430 out of 2271 proteins), and ubiquitin (Ub; right; based on 1841 out of 2578 proteins) immunoprecipitation in comparison to total neuronal lysates. Proteins were quantified by mass spectrometry and filtered based on uniqueness to the IP or fold-enrichment (>3 fold) compared to an ‘empty bead’ control. GO term overrepresentation was analyzed compared to all proteins detected in the input samples (i.e. total neuronal lysates) via a Fisher’s exact test with Benjamini-Hochberg correction (FDR <1%). **(B)** b-AP15, an inhibitor of 19S deubiquitylase activity, regulates miniature excitatory synaptic transmission, as shown by miniature excitatory postsynaptic currents (mEPSC) recorded from cultured rat cortical neurons under control conditions or with b-AP15 (5 μM, 1 hr) to inhibit 19S deubiquitylase activity. **(C)** Box plots showing the mEPSC frequency with and without b-AP15. **(D)** Box plots showing the mEPSC amplitude with and without b-AP15. Two-sample T-tests detected a significant increase in mEPSC frequency and a significant decrease in mEPSC amplitude following b-AP15 treatment (***= p<0.001; ** = p<0.01; 16 recordings from four biological replicates each).

If the 19S DUBs are enriched near synapses, what about the corresponding E3 ligases? Using synaptosome fractionation and mass spectrometry (*3*), we detected 19 E3 ligases in the mouse cortical synaptosome fraction (see Data S2) (*30*), including many well-studied Lys63-ub ligases such as Nedd4, Nedd4l, and HUWE1 (*31*, *32*). To directly visualize some of the Lys63-ub ligases at synapses, we conducted immunolabelling with antibodies against Nedd4, Nedd4I, and HUWE1 (magenta; Fig. S5) in cultured rodent neurons; we co-immunolabelled dendritic and synaptic protein markers (MAP2 and Bassoon; green). Confirming the mass spec detection in synaptosomes, we observed that all three E3 ligases were present in neuronal dendrites and dendritic spines. In addition, different E3 ligases were differentially sorted in some subcellular compartments. For example, consistent with its known role in axon development and repair (*33*), HUWE1 appeared abundant in axons (Fig. S5C) while Nedd4 and Nedd4l were less so (Fig. S5A&B). Overall, these data indicate that neuronal synapses are equipped with the enzymes for the attachment and removal of ubiquitin.

Among the many enriched synaptic targets for Lys63-ubiquitylation were proteins such as syntaxin, synaptotagmin and synaptophysin that are key players in synaptic vesicle release, suggesting a role for the dynamic Lys-63 ubiquitylation in neurotransmitter release. To test this, we measured miniature excitatory postsynaptic currents (mEPSCs) in cultured rat cortical neurons (DIV 18-21) under control conditions and following treatment with a specific small-molecule inhibitor (*34*) of 19S DUB activity, b-AP15. We first confirmed that treatment with b-AP15 (5 μM for 1 hr) induced a significant accumulation of Lys63-ubiquitylated proteins in cultured neurons (Fig. S6A), an effect that was absent following treatment with a conventional proteasome inhibitor (blocker of 20S protease activity, e.g. MG132) or an inhibitor of proteasome-dependent deubiquitylation (e.g. Capzimin) (Fig. S6A) (*35*). Upon inhibition of 19S DUB activity with b-AP15, we observed a significant increase in mEPSC frequency (Fig. 4B & C), suggesting that 19S DUB activity negatively regulates synaptic vesicle release. As the 19S also deubiquitylates Lys48-ubiquitin chains in the process of proteasomal degradation, the observed b-AP15-induced increase in mEPSC frequency could also be due to altered Lys48-ubiquitin chains on synaptic proteins. Indeed, we observed that b-AP15 also induced the accumulation of Lys48-ubiquitylation (Fig. S6A). To determine whether the altered miniature synaptic transmission was due to proteasome-dependent or proteasome-independent 19S inhibition, we measured proteasome activity upon b-AP15 inhibition in neuronal lysates. Unlike proteasome inhibitors such as bortezomib, b-AP15 did not inhibit neuronal proteasome activity (Fig. S6B-C; measured by in-gel digestion of fluorogenic substrates; see methods). This is consistent with previous studies showing that b-AP15 does not target the 19S DUB used for 26S or 30S proteasome degradation (PSMD14/Rpn11, which can be inhibited by Capzimin as shown Fig. S6A), but the other two associated DUBs (Usp14 and Uch37/Uchl5) (*28, 34*). As such, proteasome-*zw*dependent 19S deubiquitylation regulates synaptic vesicle release probability, likely by acting on the identified Lys63-ubiquitylated presynaptic proteins that are involved in vesicle exocytosis.

Within the Lys63 interactome we also detected Gria1 (See SI), an essential subunit of AMPA receptor (AMPAR) complexes that are responsible for most point-to-point excitatory synaptic transmission in the mammalian brain. Consistent with Gria1 being a target of 19S DUB activity, we found that b-AP15 also induced a significant decrease in mEPSC amplitude (Fig. 4D), which suggests an alteration in the postsynaptic sensitivity to neurotransmitters. To test directly whether free 19S DUB activity regulates dendritic AMPARs, we visualized the total dendritic AMPAR population via immunofluorescence staining (in permeabilizing conditions) under control conditions or following b-AP15 incubation. Compared to control (Fig. 5A), we observed a significant loss in the total pool of dendritic AMPARs after just 1 hr of b-AP15 treatment (Fig. 5B). Considering the ~4-day half-life of AMPARs (*22*), these data suggest that postsynaptic 19S DUB activity usually stabilizes AMPARs, protecting them from degradation. This effect was not generalized to all transmembrane receptors, as the transferrin receptor abundance was not affected by b-AP15 (Fig. S7). Because AMPARs are degraded by lysosomes (*36*), we hypothesized that postsynaptic 19S DUB activity regulates the lysosomal degradation of AMPARs. To test this, we inhibited lysosomal function using chloroquine (40 μM, 1 hr) prior to b-AP15 incubation. Indeed, lysosome inhibition rescued the loss of dendritic AMPARs induced by b-AP15 (Fig. 5A&B), suggesting that under normal circumstances the 19S DUB activity promotes AMPAR lysosomal degradation. In contrast, treatment with the proteasome inhibitor bortezomib failed to prevent the b-AP15-induced loss (Fig. 5A&B), indicating that the induced loss of AMPARs is proteasome-independent.

**Fig. 5.**
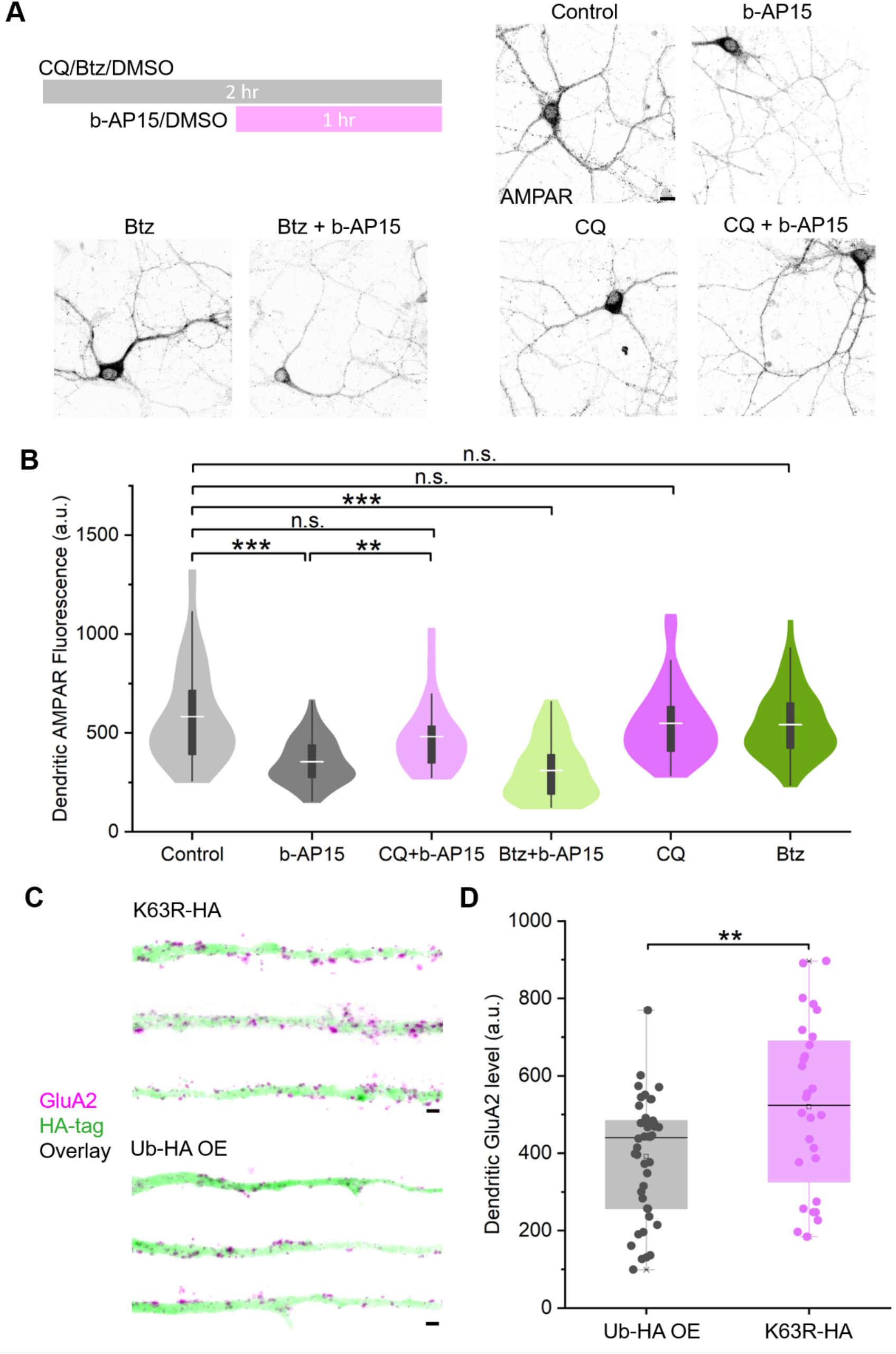
Inhibition of (proteasome-independent) 19S deubiquitylase activity regulates dendritic AMPAR abundance via Lys63 ubiquitylation. **(A**) b-AP15, a 19S deubiquitylase inhibitor, induced a loss of dendritic AMPARs that is lysosome-dependent and proteasome-independent. Upper right scheme shows the time course of the pharmacological treatment, where cultured rat cortical neurons were treated with a lysosome inhibitor (Chloroquine, CQ), a proteasome inhibitor (Bortezomib, Btz), or DMSO as control for 1 hr prior to addition of b-AP15 (or DMSO in control) for another 1 hr. The corresponding confocal micrographs immunolabelled for the total AMPAR population (anti-GluA2) were labeled following each treatment. Scale bar: 10 μm. **(B)** Violin plots showing the distribution of total dendritic AMPAR fluorescence intensity (a.u.) for each treatment show in A. Each treatment group contains >50 dendrites from > 10 neurons. ANOVA with post hoc Bonferroni test indicated significant differences between control and b-AP15, between control and CQ + b-AP15, and between b-AP15 and CQ + b-AP15 (p**<0.01, p***<0.001). **(C)** Confocal micrographs of straightened dendrites from transfected cultured neurons expressing HA-tagged K63R ubiquitin (K63R-HA) or overexpressing (OE) HA-tagged wild-type ubiquitin (Ub-HA). Samples were immunolabelled for the HA-tag (green) to identify transfected neurons, and for GluA2 (magenta) to visualize AMPAR abundance. **(D)** Box plots showing the total dendritic AMPAR fluorescence intensity (a.u.) in transfected neurons that either overexpressed wild-type ubiquitin or expressed K63R ubiquitin. A two-sample T-test detected a significant increase in the total dendritic AMPAR population in the K63R group (p**<0.01; >30 dendrites from 9 transfected neurons in each group).

The above data suggest that the free 19S particles regulate the lysosomal degradation of AMPARs via Lys63-ubiquitylation, consistent with prior knowledge of Lys63-ubiquitin signaling (*32*, *37*). To examine the importance of Lys63-ub modifications on AMPAR levels, we expressed in cultured neurons an HA-tagged ubiquitin mutant (K63R-HA) in which lysine-63 in ubiquitin is replaced with an arginine, preventing Lys63-ubiquitin chain formation. We compared the effects of K63R expression to overexpression of an HA-tagged-wild-type ubiquitin. As expected, we observed reduced total K63 ubiquitylation in neurons that overexpressed the dominant negative K63R ubiquitin (Fig. S8). We detected elevated total AMPAR levels in neurons expressing K63R, relative to control neurons which over-expressed wild-type ubiquitin (Ub-HA OE) (Fig. 5C&D). Thus, conditions that stabilize AMPAR Lys63-linkages (e.g. b-AP15 treatment) and those that destabilize them (K63R) have opposite effects on the total AMPAR pool. This directly links Lys63-ubiquitylation to AMPAR abundance, and, together with the above data, suggests that the Lys63-ubiquitylation normally destabilizes the AMPAR pool, while 19S activity opposes this destabilization.

In the above experiments, inhibition of 19S DUB activity (by b-AP15) induced a loss of AMPARs, which presumably involves accelerated endocytic sorting, a step required for surface AMPARs to enter lysosomal degradation (*32*). To test whether the 19S DUB activity normally suppresses surface AMPAR internalization, we measured the rates of AMPAR endocytosis with and without b-AP15 treatment. If 19S DUB activity normally suppresses surface AMPAR internalization, then blocking this activity should increase the internalization. We performed live-cell surface labeling of AMPARs followed by specific immunolabelling of endocytosed surface AMPARs after 1hr internalization, as described previously (*12*). Surprisingly, we found that AMPAR internalization was reduced, rather than increased, upon inhibition of the 19S DUB activity with b-AP15 (Fig. 6A-C). It thus appears that the 19S DUB activity facilitates AMPAR internalization, in contrast to previously known DUBs (e.g. Usp8) (*32*). Yet, like other DUBs (*32*), the 19S also stabilizes the total (internal + surface) AMPAR pool (Fig. 5).

**Fig. 6.**
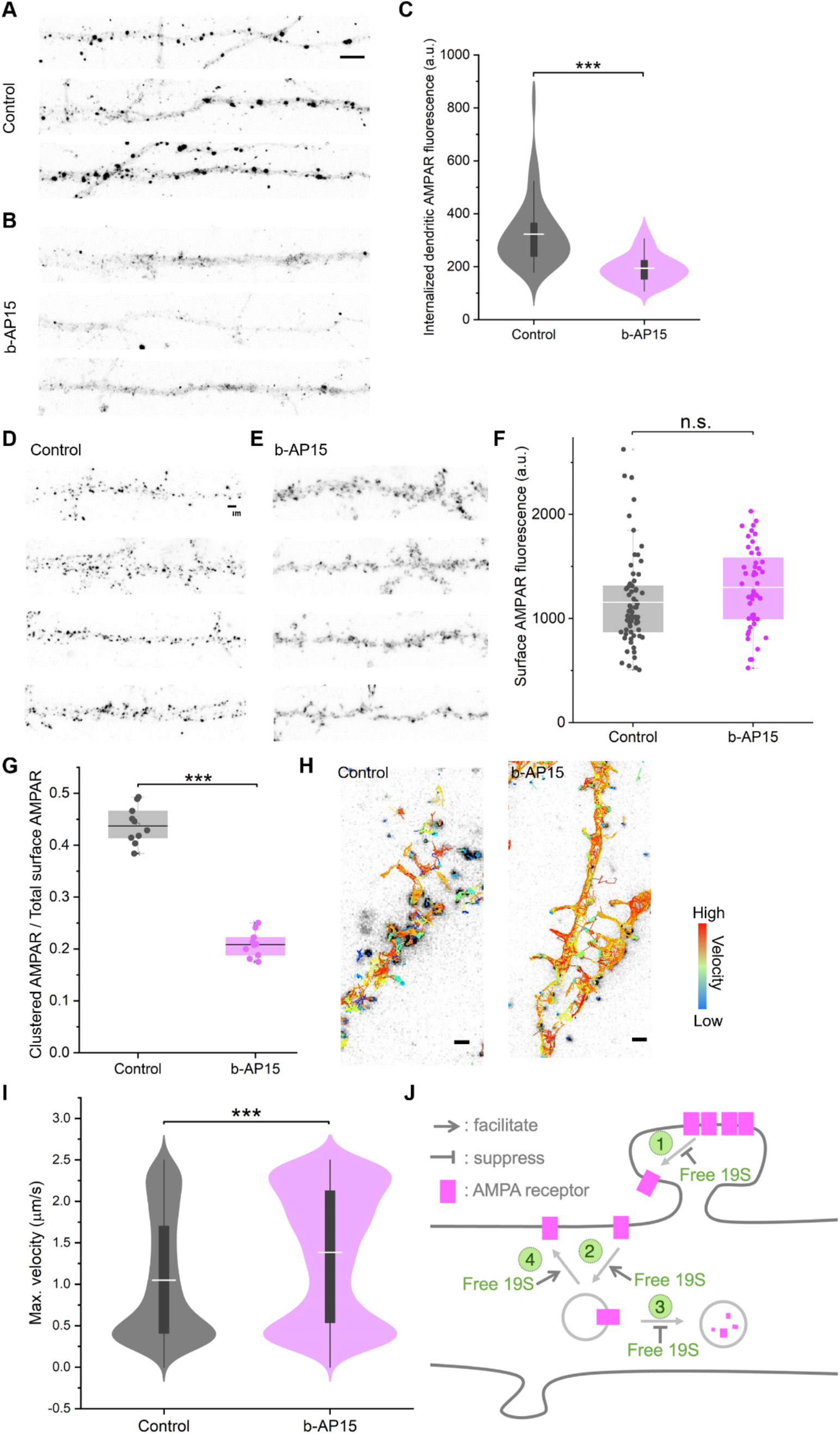
Postsynaptic 19S deubiquitylase activity maximizes synaptic availability of AMPAR. **(A) & (B)** Confocal micrographs showing internalized AMPAR in cultured rat cortical neuronal dendrites with (B) and without b-AP15 (A). Scale bar: 2 μm. **(C)** Violin plots showing the distribution of internalized AMPAR fluorescence intensity with and without b-AP15 treatment. Two-sample T-test found a significant decrease following b-AP15 treatment (p***<0.001; >50 dendrites from > 10 cells each group). **(D)&(E)** Confocal micrographs showing surface AMPARs on straightened dendrites with (E) or without b-AP15 treatment. Scale bar: 2 μm. **(F)** Box plots showing the level of surface AMPAR fluorescence in straightened neuronal dendrites with (63 dendrites from 10 neurons) or without b-AP15 treatment (48 dendrites from 10 neurons). A two-sample T test indicated no significant difference between the groups. **(G)** Box plots showing the fraction of clustered AMPARs on the cell surface with or without b-AP 15 treatment. A two-sample T-test detected a significant decrease with b-AP15 treatment. (*** = p<0.001; >10000 clusters from >10 cells in each group). **(H)** Single-particle tracking of surface AMPARs with tracking traces overlaying on the first frame. Color represents the average velocity in each tracking trace (warmer colors indicate higher velocity). 2636 tracks for control and 3397 tracks for b-AP15. Scale bar: 1 μm. **(I)** Violin plots showing the distributions of surface AMPAR mobility (max. velocity within the tracking duration) with and without b-AP15 treatment. A two-sample T-test indicated a significant increase with b-AP15 (*** =p<0.001, with 2636 tracks for control and 3397 tracks for b-AP15). **(J)** Schematic showing that free neuronal 19S DUB activity regulates the synaptic availability of AMPARs at multiple trafficking steps including 1) increasing the synaptic pool, 2) favoring the endocytosis of non-synaptic receptors, 3) reducing lysosomal degradation, and 4) facilitating exocytosis, in a concerted, proteasome-independent manner.

At first glance, the above results appeared inconsistent. We considered the possibility that differences in the number or distribution of surface AMPARs might provide an explanation. To directly test how 19S DUB activity affects surface AMPARs, we performed surface immunolabeling of AMPARs with and without the 19S DUB inhibitor b-AP15. In contrast to the b-AP15-induced reduction in total (internal + surface) dendritic AMPARs, the overall abundance of the surface AMPARs was indeed unchanged (Fig. 6D-F). Instead, the distribution of the surface AMPARs appeared altered — the receptors were less clustered and punctate following b-AP15 treatment (Fig. 6E&6G). Previous work has shown that AMPARs are mobile in the neuronal membrane but exhibit increased dwell-times at synapses, resulting in their clustering (*38*). To assess whether Lys63-ubiquitin regulates AMPAR movement, we measured the mobility of surface AMPARs via single-particle tracking in live neurons under control conditions and following treatment with b-AP15 (Fig. 6H). Inhibition of 19S DUB activity significantly elevated the mobile fraction of surface AMPARs (Fig. 6I), resulting in reduced clustering (Fig. 6G). These data suggest that Lys63-ubiquitylation of AMPARs promotes receptor mobility that can be negatively regulated (stabilized) by the free 19S de-ubiquitylase activity. Treatment with b-AP15 blocks this stabilization and results in reduced AMPAR clustering at synapses, impairing postsynaptic sensitivity to spontaneous presynaptic vesicle fusion (reduced mEPSC amplitudes; Fig.4D). Because surface AMPARs are endocytosed at sites distant from synapses (*39*), it thus appears that the free 19S subcomplex stabilizes the (presumably synaptic) clustering of surface AMPARs and, yet, facilitates their internalization once they diffuse away. Once internalized, the free 19S subcomplex then stabilizes the reserve pool of AMPARs, preventing them from degradation. In addition, we found that free 19S facilitated AMPAR exocytosis, providing an alternative to their degradation fate (Fig. S9). These results demonstrate that proteasome-independent 19S DUB activity regulates AMPARs at multiple trafficking points-all of these actions work together to maximize receptor availability at synapses (see Fig.6J for a schematic).

## Discussion

Up to 90% of the total proteome of a single neuron resides in dendrites and axons (*1*, *25*) indicating that the relative protein turnover demand at synapses is huge (*40, 41*). Local proteasomal function is thus essential for synaptic homeostasis as well as plasticity. Here we revealed a surprising differential distribution of the proteasomal catalytic and regulatory particles in neuronal cell bodies and dendrites/axons. We quantified the fractions of proteasomal particles in different proteasomal assembly states and demonstrated the presence of excess, free 19S regulatory particles near synapses. We discovered that these free 19S particles deubiquitylate important synaptic proteins that are Lys63-ubiquitylated such as AMPARs. The dynamics of Lys63 ubiquitylation and deubiquitylation potently regulate synaptic transmission in both the pre- and postsynaptic compartments. The excess of the free 19S particle present in dendrites thus represents an interesting example where a subcomplex of a large protein machine carries out a moonlighting function near synapses — in this case, as a novel ‘editor’ of a synaptic ubiquitin code.

We showed that neuronal subcellular compartments differentially localize the two main (19S regulatory and 20S catalytic) proteasomal particles. With the enrichment of 19S in neuronal processes, brain regions with denser synaptic connections such as the cortex and the neuropil of the hippocampus (*42*) also showed apparent enrichment of 19S relative to 20S (Fig. 1B). This challenges the notion that biological machines exist as a single entity throughout their life cycles (*20*). Instead, it appears that neurons have taken advantage of distinct functions of 19S particles and adapted their use to fine-tune synaptic strength. It is also possible that the free 19S may exist as a separate pool from 19S that is assembled to 20S. The question arises as to how neurons create the differential distribution of 19S and 20S particles in the process of proteasome biogenesis. Previous studies have shown that neuronal proteasomal transcripts are predominantly somatic (*43*), suggesting that proteasomal subunits are synthesized and assembled in the cell body (*15*, *44*). However, the transcripts of proteasomal subunits are not stoichiometric (*45*). To maintain stoichiometric expression (*6*, *21*), quality control pathways engage to rapidly degrade excess proteasome subunits as well as their partial assembly states that are ‘orphaned’ from 19S or 20S assembly (*45*). To survive the surveillance of quality control, proteasome subunits presumably must assemble into 19S and 20S particles rapidly post-translation. Once assembled, however, it remains unknown how 19S and 20S are sorted into different neuronal subcellular compartments (*46*).

We identified the free 19S regulatory particle as a distinct ‘editor’ of synaptic ubiquitin. For example, inhibition of proteasome-independent 19S DUB activity led to the accumulation of Lys63-ubiauitylation and resulted in abnormal presynaptic neurotransmitter release. In parallel, we identified multiple presynaptic proteins involved in synaptic vesicle exocytosis as substrates/interactors of Lys63-ubiquitylation. For example, multiple isoforms of syntaxin (stx1a, stx1b stx7, stx12), a family of membrane-anchored SNARE proteins essential for membrane fusion, were identified in our data. Consistently, syntaxin possesses Lys63-binding domains (*47*). Other important classes of synaptic proteins were also identified in the Lys63-ub interactome as potential targets regulated by free 19S, including synaptic adhesion molecules (neuroligins), ion channels, and neurotransmitter receptors. Finally, the excess free 19S near synapses may also facilitate ubiquitin recycling, as its inhibition led to accumulation of polyubiquitin chains. Consistently, previous studies have shown that accumulation of Lys48-ubiquitylated proteins upon proteasome inhibition also leads to altered synaptic transmission (*48*). As such, the free 19S synaptic DUB works in concert with the diverse local E3 ligases we and others have identified. It is also worth noting that both 19S-associated DUBs (Usp14 and Uchl5) have been implicated in neurological disorders such as ataxia and abnormal brain development (*49, 50*). So far, hundreds of E3 ligases and DUBs have been identified in mammalian cells as emerging drug targets (*14*, *51*, *52*), with many present at synapses (*27*, *30*, *32*, *53*, *54*).

We discovered an important function of the free 19S DUB activity- the concerted regulation of many aspects of the AMPAR life cycle, all of which conspire to maximize the synaptic availability of AMPARs. The free 19S proteasome directs the flow of AMPARs towards synaptic clusters at four main transit points: 1) at the plasma membrane, it facilitates the clustering of AMPARs, evident by loss of clustering upon inhibition of 19S; 2) in the extrasynaptic space, it facilitates AMPAR endocytosis; 3) for internalized AMPARs, the free 19S suppresses their degradation, 4) the free 19S also facilitates the exocytosis of AMPARs. It has been long known that 19S is sequestered at synapses during elevated synaptic activity (*5*). Till now, this has been associated only with elevated local protein degradation. Our work provides a new dimension to the role of proteasome particles in synaptic regulation— the functional states of key synaptic proteins can be regulated by free 19S via regulation of an alternative ubiquitin chain.

## Acknowledgments

We acknowledge help with transmission electron microscopy from Dr. Andre Schwartz, help with compartmentalized chambers from Claudia Fusco, help with python codes from Dr. Andreas Nold, as well as helpful discussions with Prof. Mike Heilemann.

## Funding

EMBO Long-Term Fellowship (EMBO ALTF 860-2018) (CS)

HFSP Cross-Disciplinary Fellowship (LT000737/2019-C) (CS)

Joachim Herz Stiftung Add-on Fellowship for Interdisciplinary Life Science (project number 850027) (CS)

FEBS Long-Term Fellowship (SLG)

Hessen Horizon - Marie Skłodowska-Curie-Stipendium Hessen (SLG)Max Planck Society (CS, KD, BN, SLG, PN, JL, EMS)

Advanced Investigator award from the European Research Council (grant No. 743216) (EMS)

DFG CRC 1080: Molecular and Cellular Mechanisms of Neural Homeostasis (EMS)

DFG CRC 902: Molecular Principles of RNA-based Regulation (EMS)

## Author contributions

Conceptualization: CS, EMS

Methodology: CS, KD, BN, SLG, JL, EMS

Investigation: CS, KD, BN, SLG, PN, JL, EMS

Visualization: CS, EMS

Funding acquisition: JL, EMS

Project administration: EMS

Supervision: EMS

Writing - original draft: CS, EMS

Writing - review & editing: CS, KD, BN, SLG, PN, JL, EMS

## Data and materials availability

All proteomics data associated with this manuscript have been uploaded to the PRIDE online repository (*60*). Anonymous reviewer access is available upon request. All other data are available in the main text or the supplementary materials.

## Supplementary Materials

Materials and Methods

Figs. S1 to S9

Data S1 to S2

