## Supplemental Information for "An Abundance of Free Proteasomal Regulatory (19S) Particles Regulate Neuronal Synapses Independent of the Proteasome"

**This PDF file includes:**

Materials and Methods
Figs. S1 to S9

**Other Supplementary Materials for this manuscript include the following:**

Data S1: Lys48-, Lys63- and pan-Ub pulled down proteins and GO over-representation
Data S2: E3 ligases in synaptosomes

### Materials and Methods

Cell culture and antibody staining

Dissociated rat primary cortical neuron cultures were prepared and maintained as previously reported for hippocampus neurons (*4*). In brief, cortices from postnatal day one old rat pups of either sex (RRID:RGD_734476; strain Sprague-Dawley) were dissected and dissociated by incubating with L-cysteine-papain solution at 37°C before being plated onto MatTek dishes (MatTek, Ashland, MA) previously coated with poly-D-lysine for microscopy, or 10 cm Petri dishes (MatTek, Ashland, MA) for biochemistry. Cultured cells were incubated in Neurobasal-A medium (Invitrogen, Carlsbad, CA) supplemented with B-27 (Invitrogen) and Glutamax (Invitrogen) at 37°C and 5% $CO_2$ for 18-21 days before use. Neuron cultures in compartmentalized chambers were prepared and maintained as previously described (*8*). All procedures followed national animal care guidelines and the guidelines issued by the Max Planck Society and were approved by local authorities.

Immunolabelling and Microscopy

Cultured cortical neurons in MatTek dishes were fixed in PFA-sucrose (4% paraformaldehyde; Alfa Aeser, 4 % sucrose in PBS-MC) at room temperature for 20 min, washed three times in PBS before permeabilization with 0.5 % Triton X-100 in 1 x PBS pH 7.4 for 15 min and blocking in PBS containing 4% goat serum (Gibco) for 1 hr.

For immunofluorescence staining and visualization by confocal microscopy, primary and secondary antibodies included the following:

| **Antibody** | **Source** | **Identifier** |
|---|---|---|
| **Guinea pig anti-MAP2** | Synaptic Systems | 188004 |
| **Guinea pig anti-Bassoon** | Synaptic Systems | 141004 |
| **Mouse anti-alpha7/PSMA7** | Enzo LifeSciences | MCP72 clone; PW8110 |
| **Mouse anti-Rpt5/PSMC3** | Enzo LifeSciences | TBP1-19 clone; PW8770 |
| **Rabbit anti-Rpt2/PMSC1** | ThermoFisher | HPA000872 |
| **Rabbit anti-Rpn7/PSMD6** | Abcam | ab247004 |
| **Rabbit anti-ubiquitin** | Cell signaling | 43124 |
| **Rabbit anti-Lys48 ubiquitin** | Cell Signaling | 4289 |
| **Rabbit anti-Lys63 ubiquitin** | Cell Signaling | 5621 |
| **Rabbit anti-Uchl37/Uchl5** | Abcam | ab133508 |
| **Mouse anti-HA** | Roche | 10952100 |
| **Mouse anti-GluA2** | Gift from Gouaux lab, OHSU | N/A |
| **Chicken anti-HA** | Abcam | Ab9111 |
| **Donkey anti-mouse Cy3** | Jackson ImmunoResearch | 715-165-150 |
| **Goat anti-rabbit Alexa647** | Invitrogen | A32733 |
| **Goat anti-guinea pig Alexa488** | Invitrogen | A11073 |
| **F(AB)2 Anti mouse QDOT 565** | ThermoFisher | Q11032MP |

Unless otherwise noted, overnight primary antibody staining (1:1000) was performed in PBS containing 4% goat serum (Gibco) followed by three washes with PBS, five minutes each.

Secondary antibody staining (1:1000) was performed for 1 hr in PBS, followed by three washes with PBS, five minutes each. A Zeiss LSM 880 confocal set-up was used for image acquisition.

For single molecule localization microscopy, secondary-antibody staining of proteasome particles for DNA PAINT used anti-mouse antibodies conjugated to a single-stranded DNA oligo (P1 docking) and anti-rabbit antibodies conjugated with P5 docking oligo (1:1000, custom made, sequences as previously reported) (*26*). After washing three times in PBS (5 min each), neurons were briefly fixed (5 min), and then stored in PBS at 4°C for up to three weeks until DNA-PAINT imaging commenced. For DNA-PAINT imaging, an imaging buffer containing 500 nM P1, or P5 imager oligo conjugated with Atto655 (Eurofins Genomics) in 500 mM NaCl in PBS, pH 7.4, was used as previously described (*4*). 90 nm gold fiducial markers (A1190, Nanoparz) were sparsely plated. PAINT was performed on a Leica DMi8 S system with infinity TIRF HP, Infinity Scanner, an iXon Ultra 888 EMCCD camera (Andor). A 100x oil-immersion objective (HC PL APO CORR TIRF, NA 1.47) was used in combination with a motorized TIRF illuminator. Image acquisition was performed with 638 nm excitation wavelength (150 mW) and LAS X software package. To exchange imager oligos, samples were washed three times with 500 mM NaCl in PBS before a new imaging buffer containing an alternative imager oligo was added.

Wide-field micrographs of Bassoon and MAP2 reference markers (see above Immunofluorescence labeling) were acquired for distal dendrites (at least one branch point away from the cell body) before DNA-PAINT. Super-resolution acquisition was carried out with HILO illumination and a power of 30-40 mW (determined directly after the objective and under wide-field configuration). Time-lapse datasets with 50,000 frames and 16 bit depths were acquired for each proteasome particle localization with 5 Hz frame rate and 10 MHz camera read-out bandwidth. Gain list: 2; electron multiplying gain: 100.

DNA-PAINT data were processed as previous reported (*4*). Briefly, the acquisitions were reconstructed with Picasso:Localize, a module of the Picasso software, by applying a minimal net gradient of 15000. With Picasso:Render, drift corrections were applied. Drift-corrected data were filtered using Picasso:Filter. Afterwards, raw localizations within a maximal distance of 6X measured localization precision and showing a maximum number of transient dark frames of 20 were linked together, resulting in a single, linked localization event. As shown previously (*4*), we observed minimal bleaching during imaging; in the number-of-linked-localization distribution for all the identified clusters; the smallest population of clusters was attributed as the signal of a single protein copy, with a localization precision (measured by the nearest-neighbor analysis) of 13.1 nm. To identify different assembly states of proteasomes, coincidence detection thresholds for the inter-distance between proteasome 19S and 20S clusters were set at 100 nm, taking into consideration of the localization precision, oligo-docking sites' inter-distance, image drift correction, and the actual distance between the 19S and 20S epitopes. The resulting fractions of proteasome particles in each assembly states were verified by orthogonal methods including biochemistry (Fig. 2) and existing cryo electron tomography data (*7*).

Labeling of surface AMPA receptors (AMPARs) (including for confocal microscopy and single-particle tracking), internalized and exocytosed AMPARs was conducted as previously described (*12*, *55*). Briefly, cultured rat cortical neurons were incubated with anti-GluA2 antibodies (1:100) at 37 ˚C with 5% CO2 for 5 min, followed by washing with conditioned media to remove excess free antibodies. For surface AMPAR confocal microscopy, cultured neurons were then fixed without permeabilization and stained with fluorescently tagged secondary antibodies prior to imaging. For single-particle tracking, living primary antibody-labeled cultured neurons were then live labeled with secondary antibodies tagged by quantum dots for 5 min before washing and subsequent live cell imaging (frame rate: 5 Hz; duration: 15min). Single-particle tracking data was analysed by the plugin TrackMate in Fiji. For visualization of internalized AMPARs , cultured neurons were incubated with anti-GluA2 antibodies for 20 min at 37 ˚C with 5% $CO_2$. The samples were then washed once with cold PBS-MC, incubated on ice with cold 0.5 M NaCl and 0.2% acetic acid for 4 min to strip extracellular receptor-bound antibodies, and then washed again with cold PBC-MC. The samples were then fixed, permeabilized and stained with fluorescent secondary antibodies for imaging. As control, neurons that were stained without permeabilization showed negligible signal. For visualization of exocytosed AMPARs , surface AMPARs of live cultured neurons were blocked by anti-GluA2 antibodies as well as an unlabeled secondary antibody for 5 min at 37 ˚C with 5% CO2. The excess antibodies were washed away and the cultures were allowed 1 hr in the incubator for exocytosis. Afterwards, the neurons were fixed (without permeabilization), and the newly exocytosed AMPARs were labeled with anti-GluA2 antibodies and a fluorescent secondary antibody (gt anti-ms; Alex488) for microscopy.

Biochemistry

For native protein gel electrophoresis, cortical neurons cultured in 10 cm Petri dishes were washed in ice-cold PBS buffer before they were scraped and lysed on ice using a native lysis buffer containing 50 mM Tris (pH 7.4), 1 mM dithiothreitol, 5 mM $MgCl_2$, 2 mM ATP and 250 mM sucrose. Homogenization was performed using a 1 mL syringe with a 27G needle before clarification at 4 ˚C at $10^3$ rcf for 5 min. The clarified lysate was used for native protein gel electrophoresis. The running buffer for native protein gel electrophoresis contained 500mL Tris-borate buffer (Merck; T4415), 276 mg ATP dissolved in 600 uL water and neutralized with 180 µL KOH (5 M) to pH 7, 76 mg DTT dissolved in 500 µL water, and 6.25 mL $MgCl_2$ (1 M). To condition the native gel (3-8% Criterion™ XTtris-acetate protein gel; Bio-Rad), the gel was run in running buffer for 1 hr at 50 V, room temperature. Following loading of the clarified lysates, the gel was run at 30 mA for 5-6 hrs in a cold room. The gel was then transferred onto a PVDF membrane overnight at 20 V. 4˚C before blocking and immunostaining against proteasome subunits. For in-gel proteasome activity assay, 20 mL peptide assay solution was prepared containing 1 mL Tris-HCL (pH 7.5) with 100 µL 1 M $MgCl_2$, 40 µL 0.5M ATP, and 100 µL 10 mM suc-LLVY-AMC (R&D systems). The native gel was incubated with the assay solution for 1 hr before washing with water and visualization using a Azure 280 gel imaging system (excitation, 400 nm).

For the in vitro enzymatic reaction between purified 19S and polyubiquitin chains, recombinant native cow 19S proteasome protein (ab218004) and recombinant human 26S proteasome protein expressed in HEK293 (ab218006) were purchased from Abcam. 20S proteasome protein purified from human erythrocytes (PW8720-0050) was purchased from Enzo. They were used directly at a 1:100 dilution. Recombinant human poly-ubiquitin wild-type chains (2-7) Lys63 and Lys48 were purchased from R&D Systems (UCB330) and stored as 1 mg/mL stock solution in water used in 1:100 dilution. 0.5 µL recombinant proteasome particle stock solution was incubated with 0.5 µL recombinant poly-ubiquitin chains stock solution and 19 µL buffer containing 50 mM Tris (pH 7.4), 1 mM dithiothreitol, 5 mM $MgCl_2$, 2 mM ATP and 250 mM sucrose for 2 hr at 37 ˚C before SDS-PAGE.

Immunoprecipitation

Magnetic beads conjugated with K48 TUBE (Tebu-bio; UM407M), K63 (Tebu-bio; UM404M), and control magnetic beads (UM0500M) were used following an optimized protocol from the supplier. Briefly, four dishes of cortical neurons cultured in 10 cm Petri dishes (4 Million cells per dish) were scraped, and pelleted in cold PBS before addition of 200 µL lysis buffer containing 100 mM Tris-HCl (pH 8.0), 150 mM NaCl, 5 mM EDTA, 1% NP-40, PR619 (100 µM; ubiquitin protease inhibitor; Sigma), o-phenanthroline (5 mM; DUB inhibitor; Sigma), and N-Ethylmaleimide (5 mM, DUB inhibitor; Merck). The pellets were homogenized in lysis buffer using a 1 mL syringe before centrifugation for 20 min at 4 ˚C, 16k rfc. 30 µL of clarified lysates were saved for western blotting. An 80 µL slurry of the magnetic beads was equilibrated using a TBST buffer as described by the supplier's protocol (20mM Tris-HCl pH 8, 150 mM NaCl, 0.1% Tween-20) and added into the remaining 170 µL clarified lysate for 2 hr incubation at 4 ˚C on a shaker. The beads were collected, washed, and eluted using 45 µL SDS-page loading buffer. The supernatants were saved as control. Immunoprecipitation was verified by western blotting against Lys48- and Lys63- ubiquitin. The mixture of immunoprecipitated proteins and SDS-PAGE loading buffer was prepared for mass-spectrometry.

MS sample preparation

The immunoprecipitated proteins in SDS-PAGE loading buffer were prepared for mass-spectrometry based proteomics analysis as previously reported (*56*). Briefly, samples were mixed with 2x lysis buffer (10% SDS, 100 mM TRIS, pH 7.55 with $H_3PO_4$ [supplemented with cOmplete protease inhibitor cocktail]) in a 1:1 ratio and subsequently reduced using dithiothreitol in a final concentration of 20 mM for 10 min at RT. For cysteine alkylation, the samples were incubated with iodoacetamide in a final concentration of 50 mM for 30 min at RT in the dark. Then, the samples were acidified using phosphoric acid in a final concentration of ~1.2%. Binding/wash buffer (90% methanol, 50 mM TRIS, pH 7.1 with $H_3PO_4$) was added in a 1:7 lysate-to-buffer ratio to the acidified proteins. The protein suspension was loaded onto the filter of the S-trap (size: "micro"; ProtiFi, Huntington, NY) by centrifugation for 20 s at 4,000xg in 150 µl-steps. Trapped proteins were washed with 150 µl binding/wash buffer four times. Trypsin (1µg; Promega, Madison, WI) was added in 60 µl 40 mM ammonium bicarbonate buffer. Digestion was performed overnight (~18 hrs) at RT in a humidified chamber. For peptide collection, the filter was washed in three consecutive steps by centrifugation at 4,000xg for 40 s starting with 40 µl digestion buffer and two 40 µl washes with 0.2% formic acid in MS grade water. The eluted peptides were further processed by $C_{18}$-based purification ('stage-tips') according to previously published protocols (*57*) and dried *in vacuo*.

LC-MS analysis

Dried peptide samples were reconstituted in 5% acetonitrile (ACN), 95% water and 0.1% formic acid (FA). Peptides were separated by a nano-HPLC (U3000 RSLCnano, Dionex). The samples were loaded and washed with loading buffer (2% ACN, 0.05% trifluoroacetic acid in water; 6 min; 6 µL/min) on a PepMap100 loading column (C18, L = 20 mm, ID = 75µm, 3 µm particle size, Thermo Scientific). Subsequently, peptides were separated by a gradient of water (buffer A: 100% $H_2O$ and 0.1% FA) and acetonitrile (buffer B: 80% ACN, 20% $H_2O$ and 0.1% FA) with a constant flow rate of 250 nL/min on an analytical column with an integrated emitter (C18, L= 500 mm, ID = 75 µm, 1.7 µm particle size, CoAnn Technologies; heated to 55°C). The gradient went from 4% to 48% buffer B in 90 min. All solvents were LC-MS grade and purchased from Riedel-de Häen/Honeywell (Seelze, Germany). Eluting peptides were analyzed in data-dependent acquisition mode on a Fusion Lumos mass spectrometer (ThermoFisher Scientific) coupled to the nano-HPLC by a Nanoflex source. MS1 survey scans were acquired over a scan-range of 350 to 1400 mass-to-charge ratio (m/z) in the Orbitrap detector (resolution (R) = 120k, automatic gain control (AGC) = 2e5 and maximum injection time = 50 ms). Sequence information was acquired by a "top speed" MS2 method with a fixed cycle time of 2s for the survey and after MS/MS scans. MS2 scans were generated from the most abundant precursor ions with a minimum intensity of 5e3 and charge states from 2 to 5. Selected precursors were isolated in the quadrupole using a 1.4 Da window and fragmented using higher-energy C-trap dissociation (HCD) at 30% normalized collision energy. For MS2 scans, an AGC of 1e4 and a maximum injection time of 300 ms were used. Resulting fragment ions were detected in the ion trap using the rapid scan rate. Dynamic exclusion was set to 30 s with a mass tolerance of 10 parts per million (ppm). Each sample was measured in duplicate LC-MS/MS runs.

<u>LC-MS data processing</u>

For protein identification and quantification, MS raw data were processed using MaxQuant (version 1.6.2.3; RRID: SCR_015753) with customized Andromeda parameters (*58*). For all searches, spectra were matched to a *Rattus norvegicus* sequence database (reviewed and unreviewed; downloaded from uniprot.org (RRID:SCR_004426)) and to contaminant and decoy databases. Tryptic peptides with 0 to 2 missed cleavages were considered. Precursor mass tolerance was set to 4.5 ppm and fragment ion tolerance to 0.5 Da. Carbamidomethylation of cysteine residues was selected as a fixed modification and protein N-terminal acetylation as well as methionine oxidation were selected as variable modifications. An FDR of 1% was applied at the peptide-spectrum-match (PSM) and protein level. Proteins identified by at least one unique peptide were retained for downstream analysis. Label-free quantification (LFQ) was performed by pairwise ratio determination in at least three consecutive full scans; LFQ normalization was deselected. The match-between-runs (MBR) option was enabled.

Downstream analysis of each pulled-down proteome was performed using Perseus (version 1.6.2.3; RRID: SCR_015753). First, decoy and contaminant proteins were discarded from the quantified protein groups (proteinGroups.txt). Proteins were then filtered based on uniqueness to the IP or fold-enrichment (greater than 3-fold) compared to the 'empty bead' control. GO term overrepresentation was analyzed comparing the gene names of the respective pulled down proteins to gene names of the proteins detected in input samples (i.e. total neuronal cortex lysates) via a Fisher's exact test with Benjamini-Hochberg correction (FDR <1%)

<u>Electrophysiology</u>

Cultured cortical neurons (18-21 DIV) treated with DMSO or b-AP15 (5 μM, 1 hr) were used for whole-cell recordings of miniature synaptic transmission in HEPES buffered artificial cerebrospinal fluid, as previously reported (*59*). Data were analyzed offline using Clampfit. Following a blinded manual baseline correction, mEPSC events were screened with an amplitude threshold of $\geq$ 4pA and an exponential decay.

<u>Constructs and transfection</u>

HA-wild type ubiquitin and HA-K63R ubiquitin constructs were custom-made by IDT as gBlocks gene fragments and cloned in-house. Lipofectamine 2000 (ThermoFisher) was used for transfection, following a modified protocol from the supplier. In brief, for each transfection, 2 μL lipofectamine 2000 was mixed with 75 μL neural basal medium (supplemented with Glutamax) and incubated for 5 min at room temperature. In a separate tube, 1 ug DNA was mixed with 75 μL neural basal medium (supplemented with Glutamax). The lipofectamine-containing mixture was added into the DNA-containing mixture before mixing and 20 min incubation at room temperature. The original media of cultured neurons was removed and saved before 150 µL transfection mixture was added for 1 hr incubation at 37˚C, 5% $CO_2$.Following incubation, the transfection mixture was aspirated from the cell dishes and the original media was placed back. Expression of constructs was checked after two days.

Reagents

Unless noted otherwise, all substances were molecular biology or cell culture grade and purchased from Sigma-Aldrich or Roth. MG132 (Merck) was stored as 50 mM stock solution in DMSO and used at a 1:1000 dilution. Bortezomib was stored as 2 mM stock in DMSO and used at a 1:2000 dilution. b-AP15 was purchased from Selleckchem (Catalog No. S4920; stored as 10 mM in DMSO stock solution) and used at 5 µM.

Statistics

Comparisons between two groups were conducted using two-sample t-tests. ANOVA with Bonferroni *post hoc* analysis was used for comparisons of more than 2 groups. A p value of less than 0.05 was considered significant for statistical analyses. For confocal microscopy that involved comparisons between groups, at least 10 cell replicates from 3 preparations were measured for each group. For single-molecule localization microscopy, at least 6 cell replicates from 3 preparations were measured. For biochemistry experiments, at least 3 biological replicates were used for each group. For electrophysiology experiments, 16 recordings from four biological replicates were used. Box plots include the median (50th percentile) indicated by the middle line in the box, the interquartile range (25th to 75th percentile) indicated by the top and bottom outlines of the box, the maximum and minimum without outliers (outliers defined as > 75th percentile plus 1.5 fold of the inter quartile range or < 25th percentile minus 1.5 fold of the interquartile range). Violin plots include the mean values indicated by the middle line, the interquartile range (25th to 75th percentile) indicated by the black column, the maximum and minimum without outliers indicated by the whiskers.

Fig. S1.

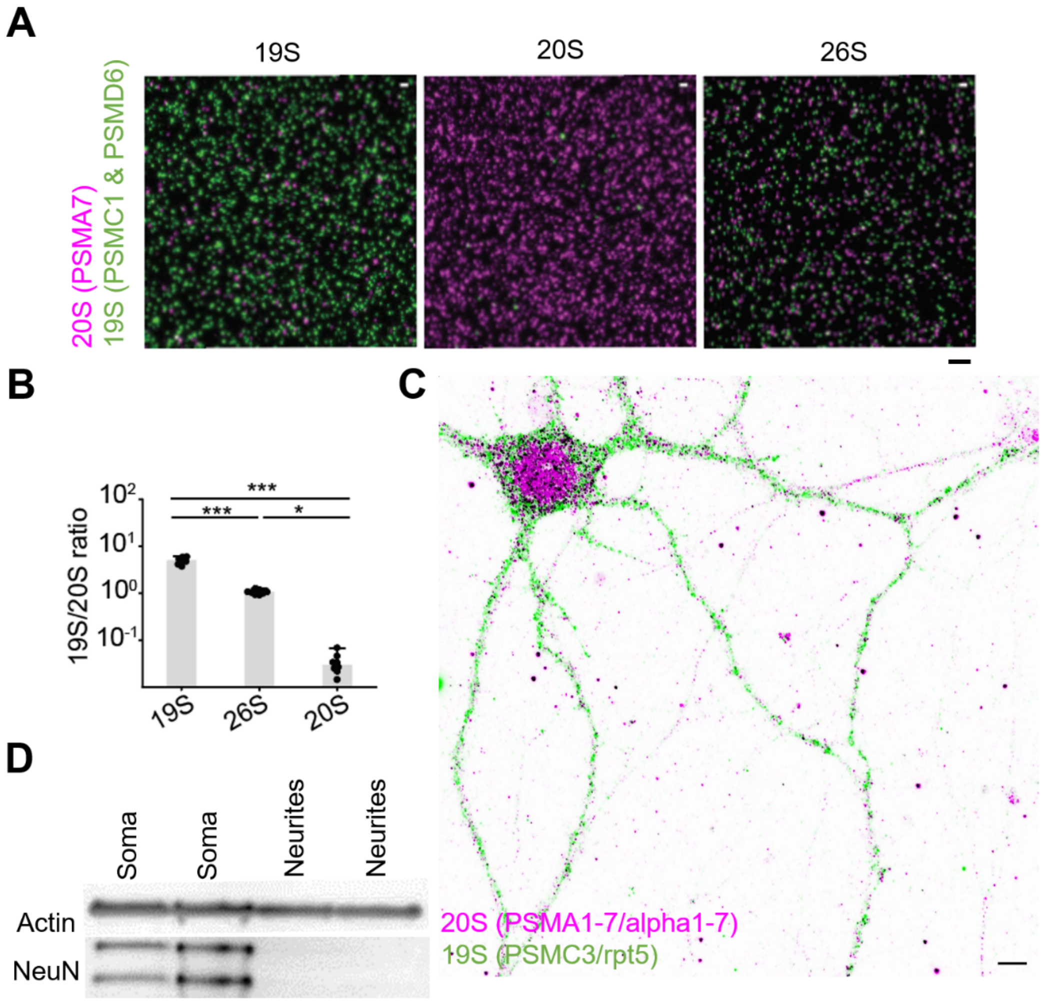


**Fig. S1. Validation of proteasome antibodies, antibody staining in neurons, and neuronal lysates from compartmentalized chambers.** (A) Confocal micrographs of purified recombinant proteasome particles 19S, 20S, and 26S plated on MatTek dishes, fixed, and immunostained with antibodies against 20S (PSMA7; magenta) and 19S (PSMC1 & PSMD6; green). Scale bar: 3 µm. (B) Bar graphs showing the ratio between 19S and 20S fluorescence puncta in MatTek dishes plated with purified 19S particles, 26S particles, and 20S particles. ANOVA with *post hoc* Bonferroni test found significant differences between the three (8 replicates each group, with 26S group averaged ~1). *=p<0.05; ***=p<0.001. (C) Confocal micrograph of a cultured rat hippocampal neuron immunostained against 20S (magenta) and 19S (green) targeting alternative epitopes. Overlay is in black. Scale bar: 5 µm. (D) Western blots showing depletion of a nuclear protein staining (NeuN) in neurite-enriched lysates harvested from compartmentalized chambers in comparison to soma-enriched lysates.

**Fig. S2.**

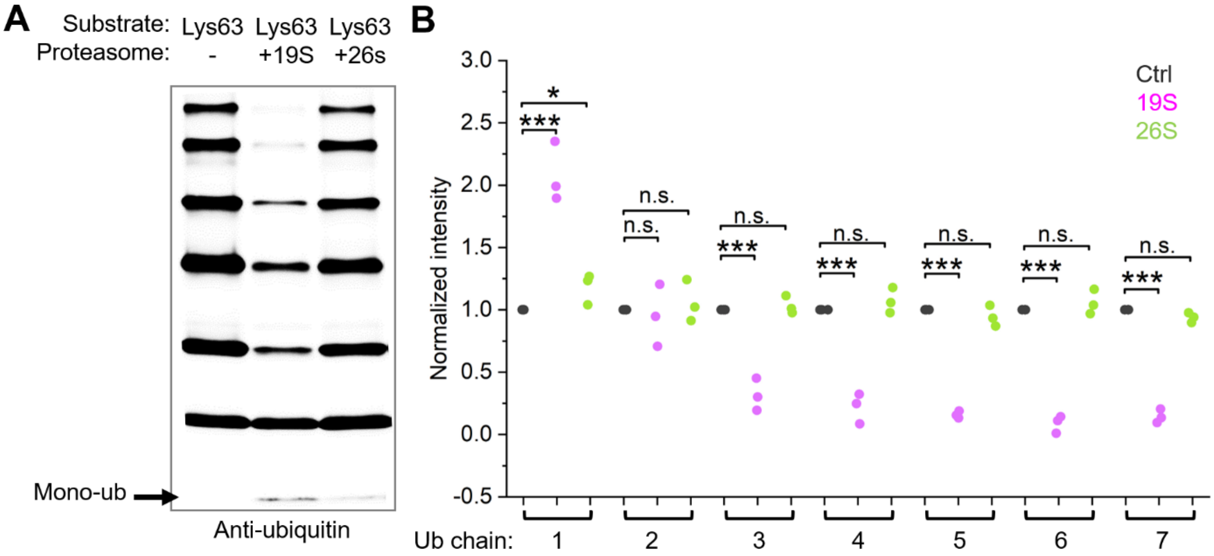


**Fig. S2. Purified recombinant 19S degrades Lys-63 polyubiquitin chains.** (A) Purified recombinant 19S and 26S proteasome particles were incubated with Lys63-polyubiquitin chains for 1 hr (see methods) before SDS-PAGE and immunolabelling with an anti-ubiquitin antibody. (B) The scatter plots show the intensities of different ubiquitin chain lengths (1-7) in each lane (three replicates each; normalized by control). ANOVA with Bonferroni post hoc analysis indicated a significant difference between the control and 19S for most chain lengths, and no significant difference between the control and 26S lane for most chain lengths (***= p<0.001; *= p <0.05).

Fig. S3.

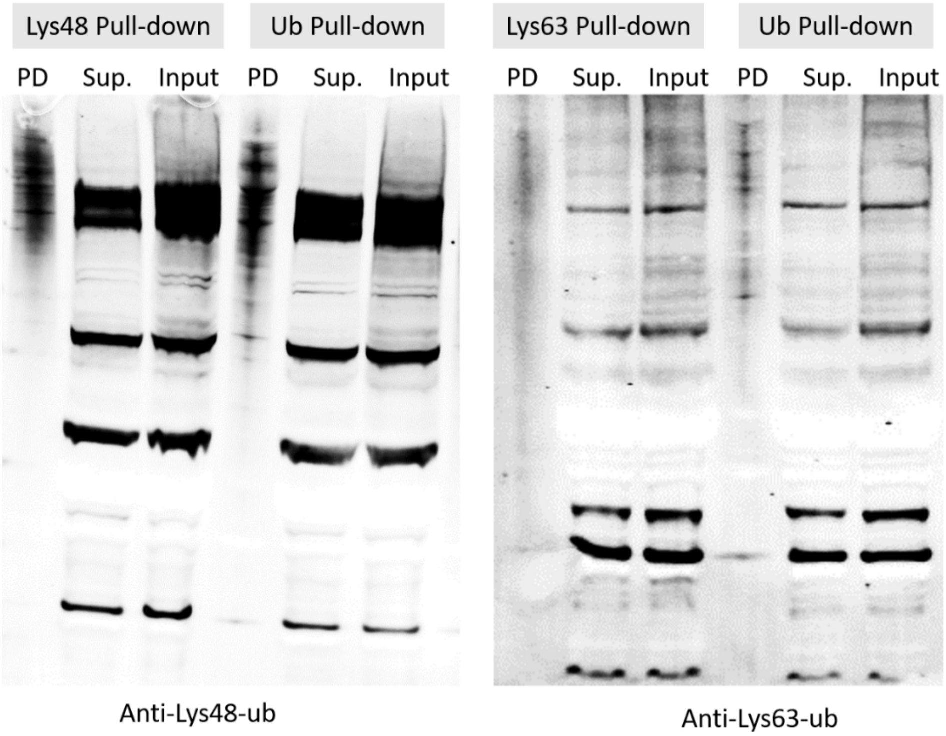


**Fig. S3. Immunoprecipitation of Lys48-ub, Lys63-ub, and ubiquitin (Ub) bound neuronal proteins.** Blots on the left were immunolabelled with an anti-Lys48-ub antibody while blots on the right were immunolabelled with an anti-Lys63-ub antibody. Proteins that were immunoprecipitated (pulled-down, PD) in Lys48-ub, Lys63-ub, and ubiquitin (Ub) immunoprecipitations were compared to proteins in the supernatant (Sup.) and to the input of the respective immunoprecipitation. Multiple clearly-defined, intense bands in the Supernatant And input lanes were off-target background recognized by the anti-lys48 and -lys63 antibodies and were absent in the pull-down lanes.

**Fig. S4.**

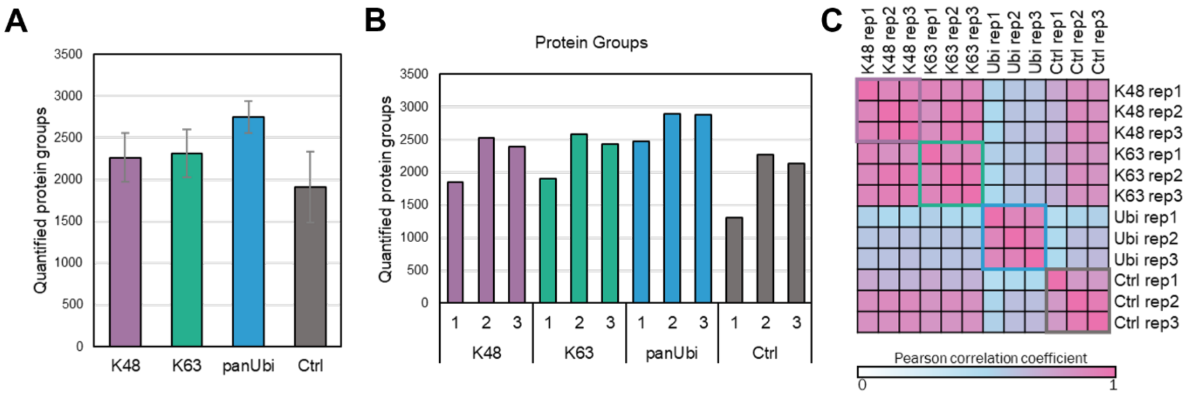


**Fig. S4. Mass spectrometry of neuronal proteins in the pull-down samples of Lys48-ub, Lys63-ub, and ubiquitin (pan-Ubi) immunoprecipitations.** (A) shows the total number of proteins identified in Lys48 (K48)-ub, Lys63 (K63), ubiquitin with unspecified linkages (panUbi), and control (naked beads) pull-down samples. (B) shows the number of proteins identified in each biological replicate of each group. (C) shows the correlation coefficients amongst the biological replicates.

**Fig. S5.**

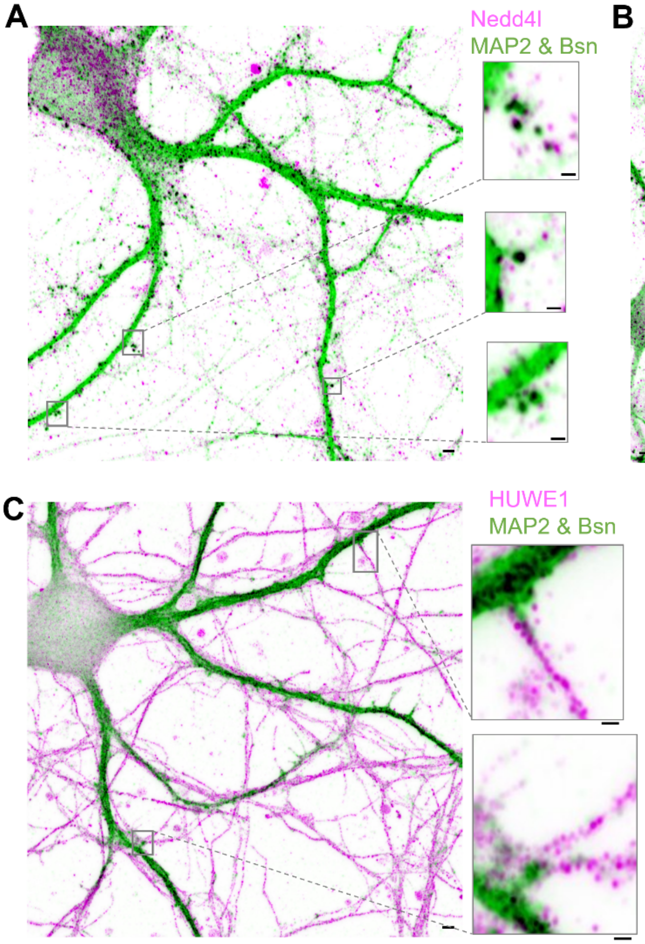


**Fig. S5. Diverse E3 ligases are present at neuronal dendrites, axons, and dendritic spines**. **(A)-(C)** show confocal micrographs of three synaptosome-enriched E3 ligases (magenta) immunostained in cultured cortical neurons together with dendritic and synaptic markers, MAP2 and Bassoon (Bsn), in green, and their overlay in black. Scale bars: 2 µm. Right inset of each confocal micrograph shows enlarged examples of dendritic spines present with respective E3 ligases. Scale bars of inset: 1 µm.

**Fig. S6.**

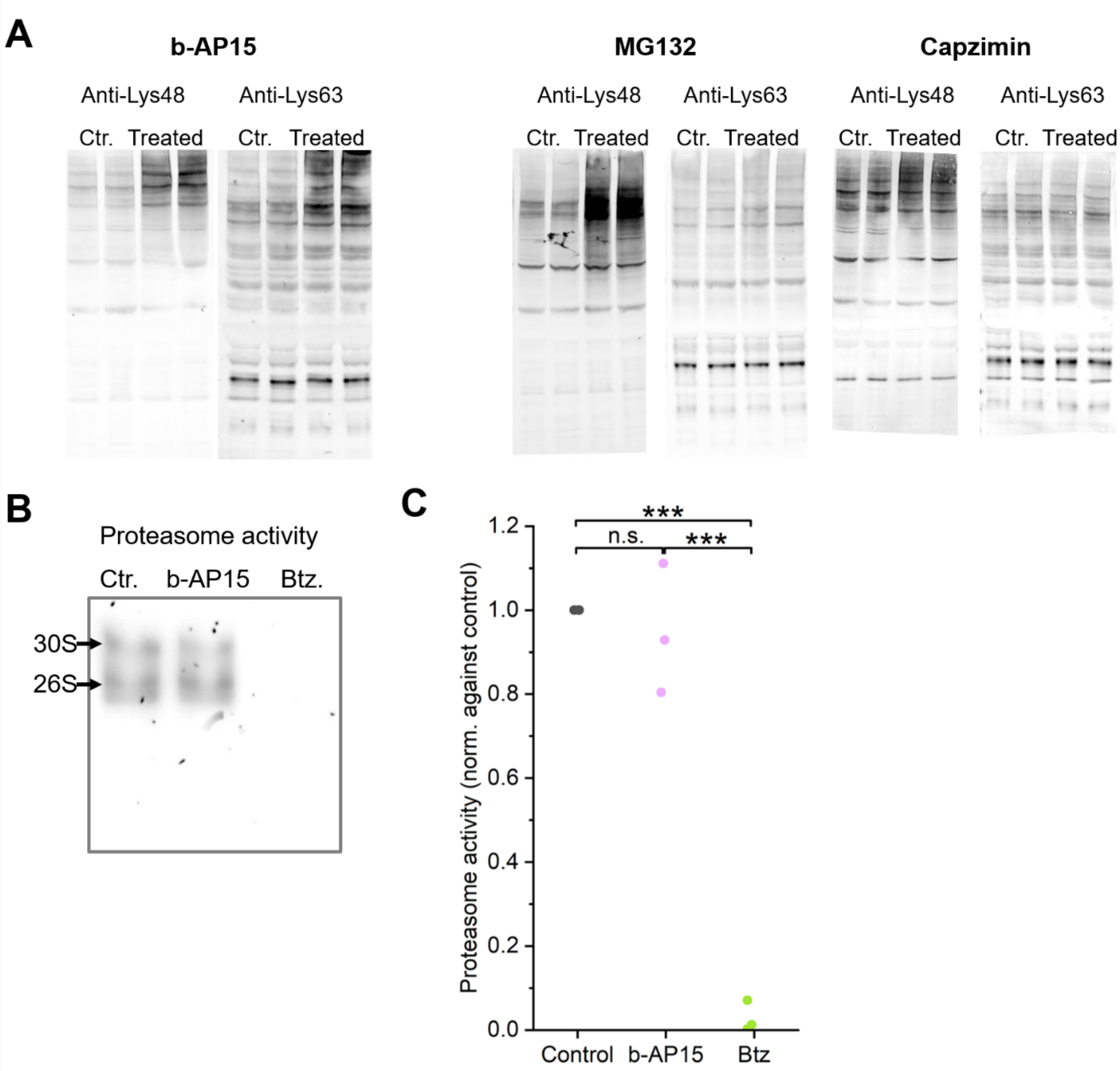


**Fig. S6. b-AP15 inhibits proteasome-independent deubiquitylation activity. (A)** SDS-page gel of neuronal lysates immunolabelled with an anti-Lys48-ubiquitin antibody or an anti-Lys63-ubiquitin antibody after *in vitro* treatment with b-AP15 (5 µM, 1 hr), MG132 (50 µM, 2 hr), or Capzimin (5µM, 15 hr). **(B)** Proteasome activity of neuronal lysates was probed by in-gel incubation with the fluorogenic peptide suc-LLVY-AMC following native protein gel electrophoresis (see methods). The two arrows highlight the 30S (high) and 26S (low) proteasomes. **(C)** Scatter plots showing proteasome activity measured by fluorescence intensity in B (normalized to control). ANOVA with post hoc Bonferroni test found no significant differences between control and b-AP15 treated groups. The established inhibitor Bortezomib (2 µM, 2 hr) significantly inhibited proteasome activity (*** = $p<0.001$).

**Fig. S7.**

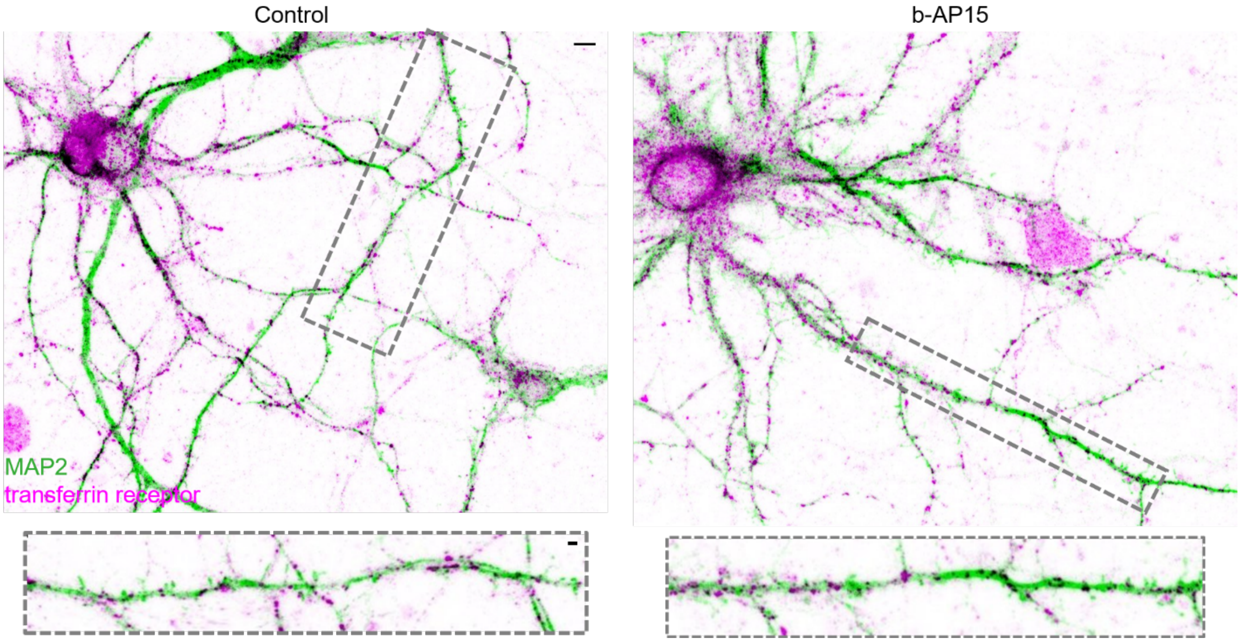


**Fig. S7. Transferrin receptor levels are not affected by b-AP15 treatment.** Confocal micrographs showing cultured rat cortical neurons treated with DMSO (control; left) or b-AP15 (5 µM; 1 hr) and immunolabelled with an anti-MAP2 antibody (green) and an anti-transferrin receptor antibody (magenta). Scale bars: 5 and 2 µm in upper and lower image, respectively.

**Fig. S8.**

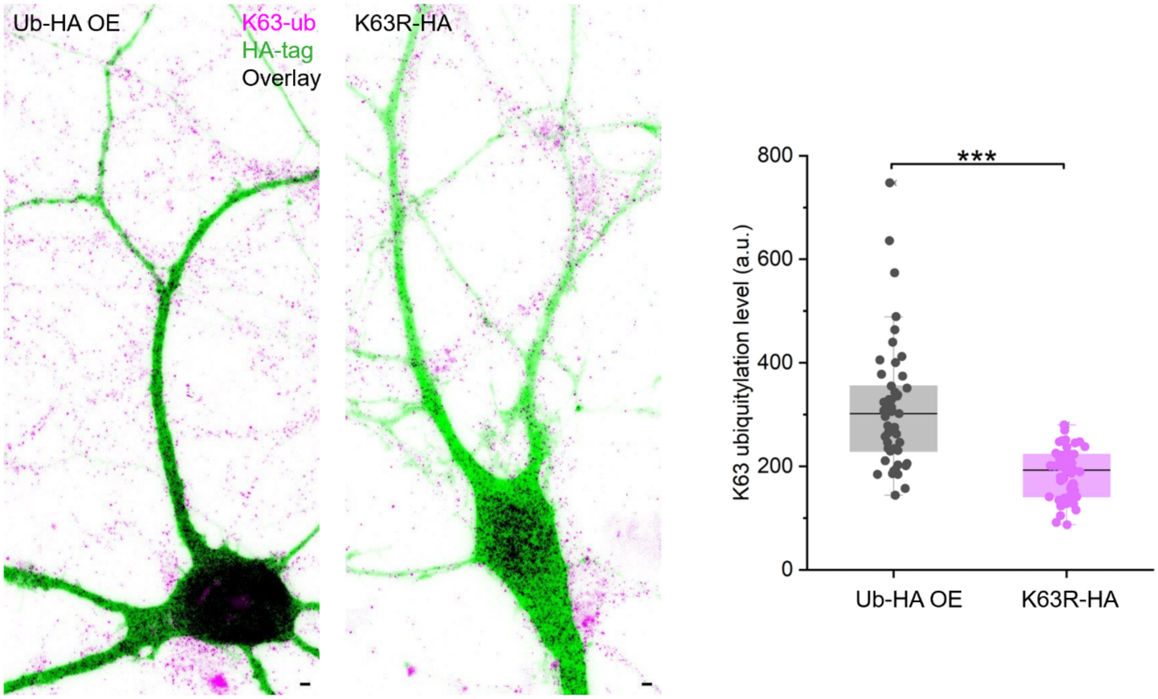


**Fig. S8. Overexpression of K63R ubiquitin reduced Lys63-ubiquitylation.** Confocal micrographs of transfected neurons overexpressing an HA-tagged wild-type ubiquitin (Ub-OE) or HA-tagged K63R ubiquitin (K63R) immunolabelled within an anti-HA antibody (green) and an anti-Lys63-ubiquitin antibody (magenta). The overlay between the two is in black. Box plots on the right show the Lys-63 ubiquitylation level in >48 dendrites from 10 cells in each group. A two-sample t-test indicate a significant decrease in transfected neurons that overexpressed K63R (***= $p<0.001$). Scale bar: 2 µm.

**Fig. S9.**

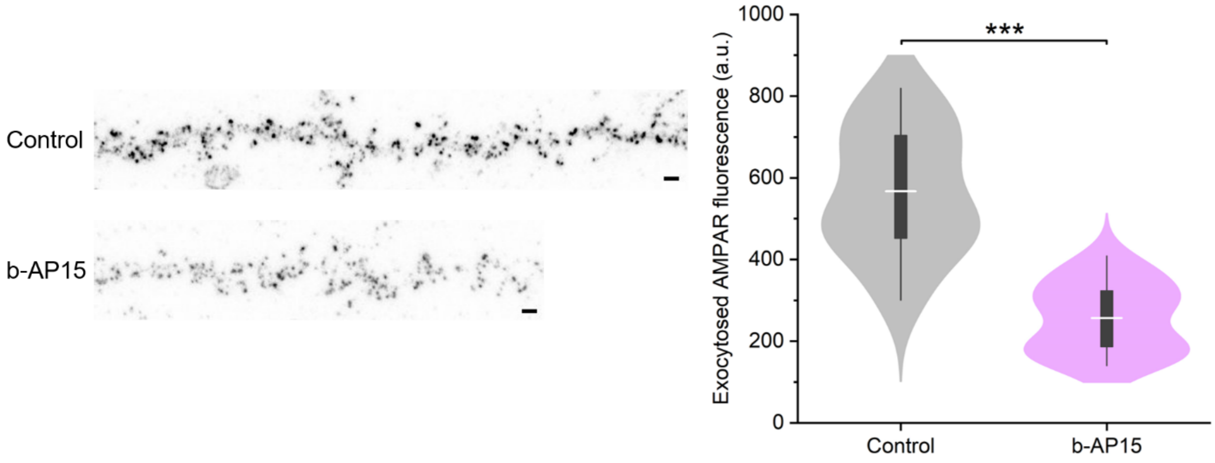


**Fig. S9 Free 19S facilitates exocytosis of AMPA receptors.** Confocal micrographs of surface AMPA receptors that were exocytosed within 1 hr before fixation and immunostaining. Shown are straightened (via image processing) dendrites from cultured neurons treated either with 19S DUB activity inhibitor, b-AP15, or DMSO as control. Right violin plots showed the exocytosed AMPA receptor fluorescence in control and b-AP15 treated groups. Two-sample T test found a significant decrease following b-AP15 treatment. (>20 dendrites from 8 cells for each group; ***= $p<0.001$). Scale bar: 1 µm

**Data S1. (separate file)**

Lys48-, Lys63- and pan-Ub pulled down proteins and GO over-representation

**Data S2. (separate file)**

E3 ligases in synaptosomes
